# Voclosporin Preserves Mitochondrial Function Compared With Cyclosporine A in Perfused Human Proximal Tubule Microphysiological Systems

**DOI:** 10.64898/2026.07.07.737071

**Authors:** Kayenat S. Aryeh, Yik Pui Tsang, Eric W. Hsu, Catherine K. Yeung, James MacDonald, Theo K. Bammler, Jonathan Himmelfarb, Linda M. Rehaume, Edward J. Kelly

## Abstract

**Key Points:**

- Perfused human kidney MPS revealed CsA-associated sublethal tubular stress that was not detected by conventional 2D viability assays or by KIM-1 release in 3D MPS.
- At matched exposure, VCS preserved mitochondria and activated ER chaperones and iron detoxification, with no p21 arrest compared to CsA.
- Mechanistic separation supports VCS’s nephroprotection potential and early mechanism-based biomarkers to guide CNI choice.

Graphical Abstract

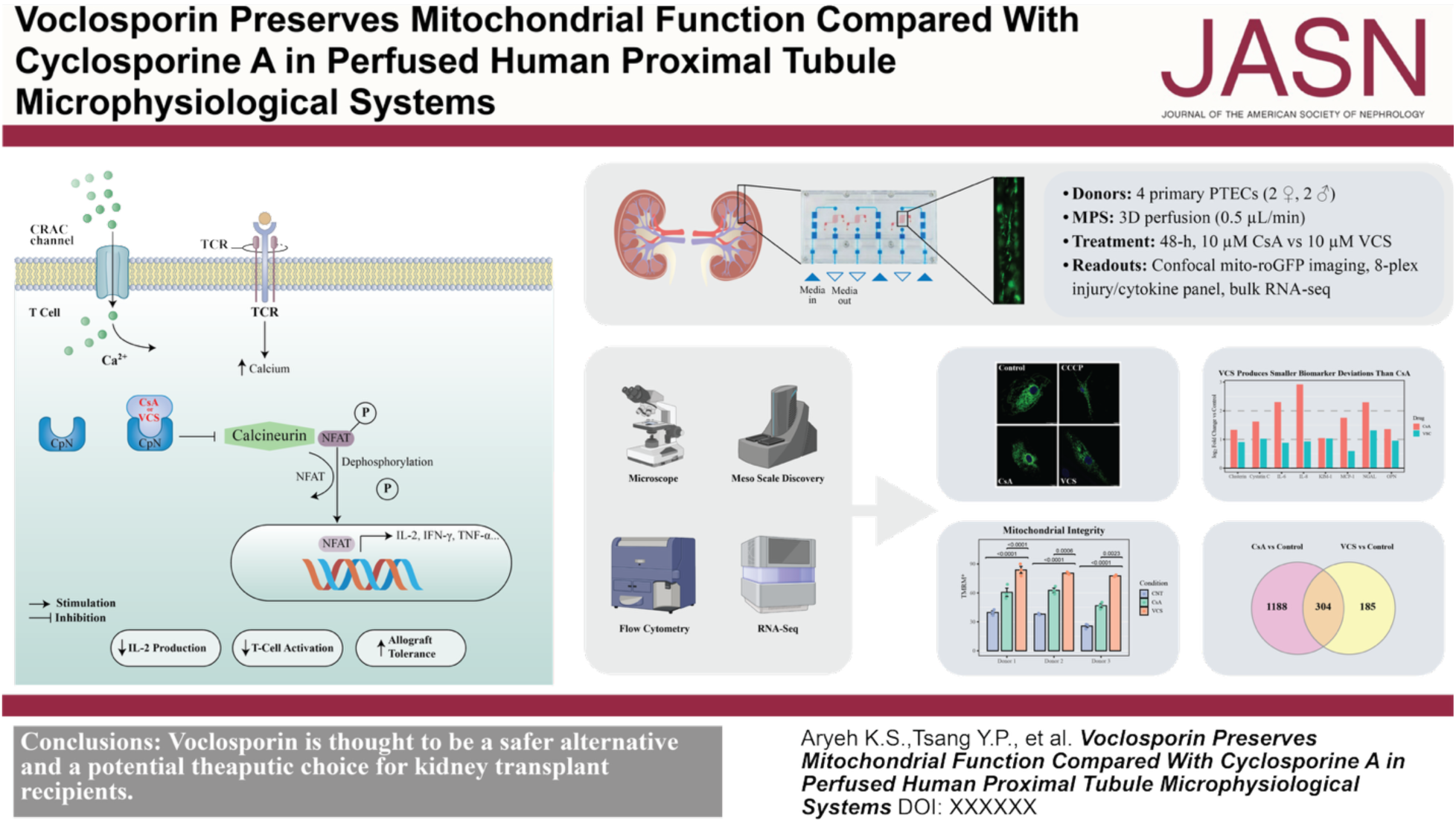

**Background:** Calcineurin inhibitors (CNIs) are indispensable for transplantation immunosuppression, yet cyclosporine A (CsA) produces renal toxicity. Voclosporin (VCS), a CsA analog, is proposed to be less nephrotoxic, but mechanisms remain unclear.

**Methods:** Primary human proximal tubule epithelial cells (PTECs) were exposed to CsA or VCS in 2D monolayers and perfused 3D kidney microphysiological system (MPS). Viability was assessed in 2D cultures by MTS, mitochondrial membrane potential (ΔΨm) by TMRM flow cytometry, and soluble injury and inflammatory biomarkers in MPS effluents by ELISA and MSD multiplex assays. RNA sequencing of 3D-cultured PTECs was used to identify differentially expressed genes and pathways.

**Results:** In 2D PTECs, neither drug reduced viability. In 3D MPS effluents, KIM-1 did not distinguish CsA from VCS, whereas the MSD biomarker panel showed larger aggregate deviation with CsA. Confocal tomography showed CsA-associated mitochondrial fragmentation, whereas VCS preserved reticular mitochondrial architecture. TMRM flow cytometry showed a treatment-dependent difference in TMRM-positive cells, with VCS yielding the highest TMRM-positive fraction and exceeding CsA, supporting preservation of ΔΨm relative to CsA. RNA-seq identified 1188 CsA-specific and 185 VCS-specific differentially expressed genes, with 304 shared. Pathway analysis indicated CsA enrichment of unfolded protein response (UPR) and endoplasmic reticulum (ER) stress, p21-associated G_2_/M checkpoint arrest, and transcriptional signatures consistent with ferroptosis priming, while VCS mainly induced ER chaperone and ER-associated degradation gene programs without activating canonical UPR sensors and showed limited cell-cycle suppression.

**Conclusions:** A physiologically relevant 3D kidney MPS revealed sublethal tubular stress from CsA that is masked in 2D culture, including mitochondrial depolarization, proteostatic stress, and ferroptosis priming. At matched exposure, VCS preserved mitochondrial function and proteostasis while eliciting a narrower, adaptive ER quality control response. These data support VCS as a nephron-sparing immunosuppressant and 3D MPS as a mechanism-based platform for evaluating renal safety of drugs and nominating early sub-lethal tubular injury biomarkers.

## Introduction

Calcineurin inhibitors (CNIs) are essential for preventing acute rejection after solid-organ transplantation, yet nephrotoxicity produced by first-generation agents such as cyclosporine A (CsA) compromises long-term graft survival^1–3^. Chronic exposure disrupts mitochondrial function, promotes oxidative stress, and drives tubulointerstitial fibrosis^3–6^. Voclosporin (VCS), a single-amino-acid analog of CsA, has a more predictable exposure profile, including smaller peak-to-trough fluctuation and near dose-proportional systemic exposure across clinically relevant doses. In phase 3 trials, VCS was associated with smaller declines in eGFR in phase 3 trials^4–6^, however, the cellular basis of this renal advantage remains undefined.

Traditional two-dimensional (2D) culture systems remain widely used for early drug screening because of their simplicity, throughput, and cost^7^. However, they fail to recapitulate the architectural and physiological complexity of the renal microenvironment and can obscure early organellar stress. *In vivo*, proximal tubule epithelial cells (PTECs) experience luminal shear stress, three-dimensional (3D) tubular architecture, dynamic cell-cell and cell-matrix interactions, and gradient-based solute delivery, all of which are critical to sustain proper tubular function and cellular homeostasis^8,9^. The absence of these factors in 2D systems can mask subtle mitochondrial and ER stress responses that precede overt damage. By contrast, perfused 3D kidney microphysiological systems (MPS) restore key microenvironmental features and enable detection of sub-lethal cell toxicities and stress phenotypes not evident in static monolayers^10^.

Mitochondrial health is a central determinant of PTEC viability and resilience. Loss of mitochondrial membrane potential (ΔΨm) and redox imbalance can propagate to ER proteostasis pathways and sensitize cells to iron-dependent lipid peroxidation^11–13^. Given VCS’s distinct pharmacology, we hypothesized that VCS would evoke a more adaptive stress pathway than CsA, which spares mitochondrial bioenergetics and proteostasis pathways that govern tubular recovery. Here, we used a human perfused 3D kidney MPS to compare CsA and VCS at clinically relevant concentrations. We integrate high-resolution confocal imaging of the mitochondrial network, tetramethylrhodamine-based ΔΨm assays, soluble injury biomarkers and cytokines, and bulk RNA sequencing to map the trajectory from initial organellar stress to transcriptional remodeling across primary PTECs from multiple donors. We further relate these molecular signatures to conventional injury readouts to identify gaps in current biomarker sensitivity.

## Methods

Detailed methods, reagent list, device schematics, and computational pipelines are provided in the Supplemental Material.

### PTEC Isolation, Culture, and Seeding in 3D Kidney MPS

Primary human PTECs (2 female and 2 male donors; Table S1 for donor demographics) were isolated from non-transplantable, normal adult donor kidneys (GFR > 60 mL/min) obtained via a U.S. Organ Procurement Organization (Novabiosis, Durham, NC, USA). The kidneys were preserved in sterile conditions at 4°C with less than 36 hours of cold ischemic time. Kidney cortex tissue was enzymatically digested with collagenase type IV (0.75 mg/mL) and dispase (0.75 mg/mL) at 37°C for 1 hour, filtered through a 100 μm strainer, and cultured in PTEC medium (DMEM/F12 media with low glucose (1 g/L), supplemented with 10 mM HEPES, 14 mM sodium bicarbonate, 1.72 µM insulin, 68 nM transferrin, 38.7 nM sodium selenite, 100 units/mL of penicillin, 100 μg/mL of streptomycin, and 0.25 μg/mL of Amphotericin B) as previously described^9,14^. PTECs were grown to confluence and used either as 2D monolayers or in a 3D perfused kidney MPS. 3D MPS devices (Nortis, Bothell, WA) contained a collagen I (rat tail, 6 mg/mL) hydrogel with a microchannel coated with mouse collagen IV (5 µg/mL) (Figure 1). PTECs were seeded at 15 to 20 × 10⁶ cells/mL, allowed to adhere for 24 hours, and then perfused continuously with PTEC medium at 0.5 µL/min. Lumen formation, barrier integrity, and viability were monitored microscopically for 1 to 2 weeks before drug treatment.

**Figure 1.**
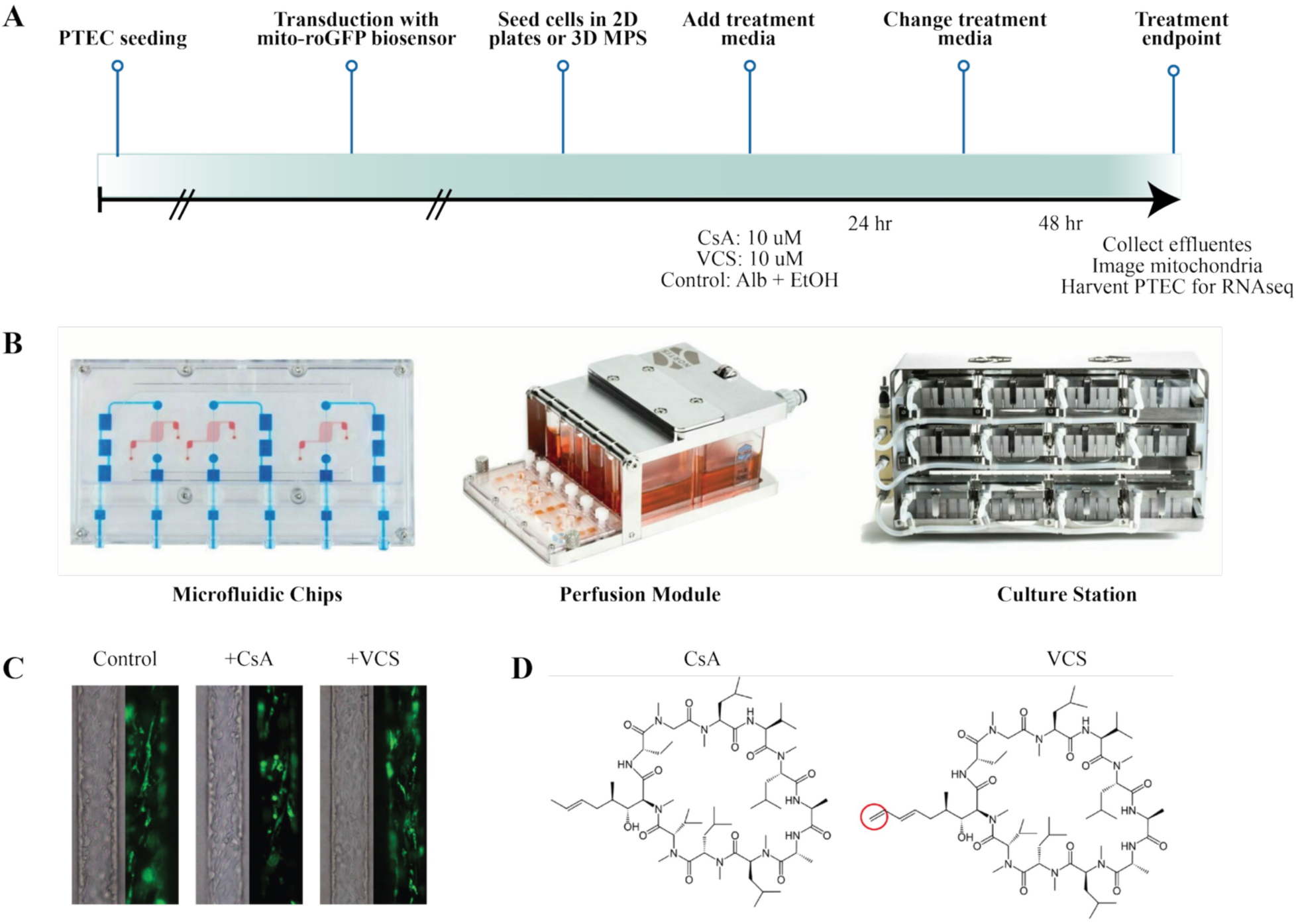
Experimental workflow, perfused proximal tubule MPS, and study compounds. (A) Primary human PTECs were expanded, transduced with a mitochondrial matrix-targeted mito-roGFP (COX8–GFP) reporter, seeded as 2D monolayers or into 3D kidney MPS, and exposed for 48 hours to CsA or VCS at 10 µM. Media contained human serum albumin at 100 µg/mL. Vehicle was ethanol-matched (0.1% v/v). Effluents, RNA, and high-resolution mitochondrial images were collected at the 48-hour endpoint. (B) Microfluidic chips were embedded in collagen and connected to a recirculating perfusion module and culture station. Channels were formed in collagen I (6 mg/mL) and coated with collagen IV (5 µg/mL). Flow rate was set to 0.5 µL/min. (C) Representative confocal micrographs of mito-roGFP**–**expressing PTECs after vehicle, CsA (10 µM), or VCS (10 µM) treatments for 48 hours. Green: mitochondrial matrix; tubule diameter of 120 μm. (D) Chemical structures of CsA and VCS. The methylene extension unique to VCS is highlighted in red.

### Drug Treatments

CsA and VCS were purchased from ChemCruz (Dallas, TX, USA). Stocks were prepared in ethanol, and vehicle control treatments were normalized to 0.1% v/v of ethanol. Cells in 2D and 3D platforms were exposed to clinically relevant concentrations of CsA or VCS (0, 1, 5, 10 µM) for the acute 48 hour experiments. Where indicated, 3D MPS cultures were maintained for 14 days to assess longer-term KIM-1 release, reported in Figure S1. Media were refreshed every 24 to 48 hours. To approximate protein binding, treatments contained human serum albumin at 100 µg/mL.

### Assessment of Injury and Mitochondrial Function in CNI-treated PTECs

PTEC viability in 2D monolayers was measured by MTS (3-(4,5-dimethylthiazol-2-yl)-5-(3-carboxymethoxyphenyl)-2-(4-sulfophenyl)-2H-tetrazolium) colorimetric assay (Abcam) according to the manufacturer’s protocol. For the assessment and imaging of the mitochondrial network, PTECs were transduced with a mitochondrial-targeted COX8-GFP reporter (lentivirus, Multiplicity of Infection = 10:1). Confocal images were acquired on a Nikon A1R. For the measurement of mitochondrial membrane potential, ΔΨm was quantified using tetramethylrhodamine methyl ester (TMRM) staining with flow cytometry (BD Symphony A3). Carbonyl cyanide m-chlorophenyl hydrazone (CCCP, 1 µM) was used as a positive depolarization control. Cytometry data were processed in FlowJo X. For the quantification of kidney injury biomarkers and cytokines from 3D MPS effluents, samples were collected at 24-hour intervals and stored at −80°C. Kidney injury markers were measured by DuoSet ELISA kits (R&D Systems) or multiplexed electrochemiluminescence assays (Meso Scale Discovery, MSD) (Meso Scale Daignostics, Rockville, Maryland, USA). Pro-inflammatory cytokines (e.g., IL-6, IL-8) were quantified by validated MSD multiplex assays.

### RNA Sequencing and Bioinformatics

Total RNA from 3D-cultured PTECs was isolated using RNeasy Micro Kit (Qiagen) and quality-checked on an Agilent Bioanalyzer (RNA integrity number > 9). Libraries were prepared using SMARTer Stranded RNA Library Preparation kit (Takara Bio) and sequenced on Illumina NovaSeq 6000 and NovaSeq Plus X platforms. Raw data were assessed using fastqc and then aligned against the GRCh38 genome using the HiSat2 aligner.^15^ Counts per gene were imported into R using the Bioconductor Rsubread package.^16^ Differences between treated and control groups were assessed using the Bioconductor limma package and its limma-voom pipeline.^17^ Due to incomplete pairing for each subject, the data were analyzed using a linear mixed model with blocking on subject, or individually by subject. Genes were selected using a Benjamini-Hochberg false discovery rate (FDR)<0.05. Pathway enrichment and analyses were conducted via KEGG and iPathwayGuide (Advaita Corporation, Ann Arbor, MI), with additional corroboration using ENRICHR and BioJupies databases.

### Statistical Analysis

Statistical analyses were performed in RStudio (version 2024.04.2) unless stated otherwise. Data are presented as mean ± standard deviation (S.D.) or standard error (SEM) as indicated in figure legends. Group comparisons used one-way or two-way ANOVA with Dunnett’s or Šidák’s post-hoc corrections for multiple comparisons. Pairwise comparisons were evaluated using two-tailed t-tests where appropriate. A P value < 0.05 was considered statistically significant. Biological replicate numbers (donors) and technical replicates (channels (3D) or wells (2D)) are specified in each figure legend.

## Results

### CsA and VCS preserve PTEC viability and KIM-1 but diverge in multiplex biomarkers

MTS assays in 2D monolayers showed no cytotoxicity across 0.01–10 µM. Viability remained ≥ 95% in PTECs from three donors and ≥ 88% in the fourth (Figure 2A). In the perfused 3D MPS, KIM-1 remained at baseline during the 48-hour study (Figure 2B) and throughout 14 days of exposure (Figure S1). Accordingly, we quantified viability only in 2D, where MTS readouts are reliable, and measured KIM-1 only in 3D MPS, where low basal KIM-1 and perfusate sampling provide a sensitive, non-destructive injury readout. In contrast, a multiplex panel of FDA-qualified kidney injury biomarkers plus IL-6, IL-8, and MCP-1 revealed clear treatment differences in MPS effluents (Figure 3). Because donor availability and computable matched fold-change values varied across endpoints, we plotted donor-level values and analyzed biomarker-level paired summaries using the absolute log₂ fold change from matched vehicle (Table S2). CsA generally produced larger absolute deviation from vehicle than VCS, although OPN at 24 h was an exception. After averaging the 24 h and 48 h endpoints within each biomarker, CsA produced greater aggregate absolute deviations than VCS (mean paired differences, CsA – VCS = 0.72 log₂ units; 95% CI, 0.37-1.07; paired t-test; P = 0.0017; Wilcoxon signed-rank P = 0.0078; Table S3). Endpoint-specific paired analyses showed a near-significant trend at 24 h (mean difference = 0.61 log₂ units; 95% CI, −0.01-1.23 paired t test, P = 0.0526) and a significant difference at 48 h (mean difference = 0.83 log₂ units; 95% CI, 0.32-1.34; paired t test, P = 0.0064; Table S3).

**Figure 2.**
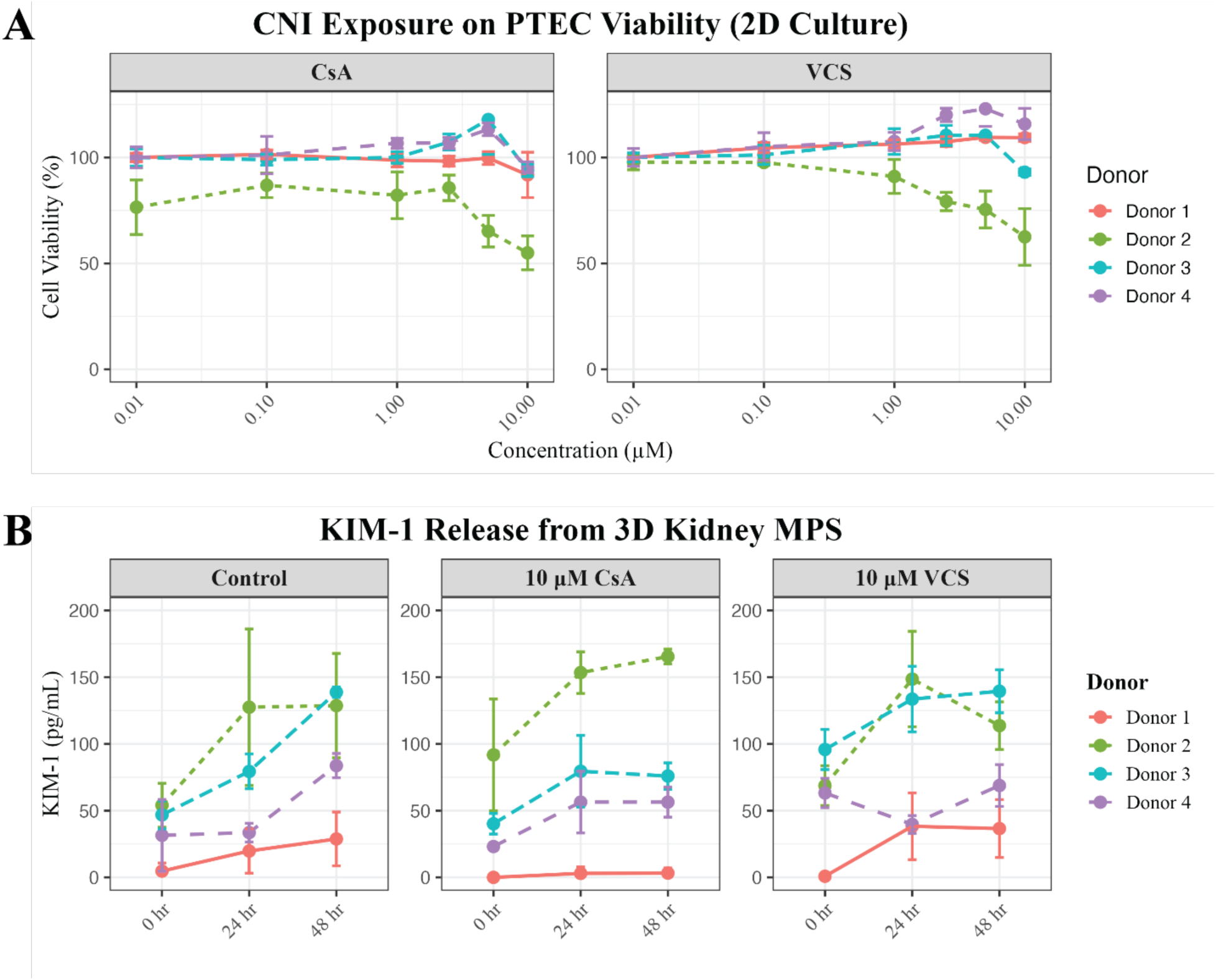
Viability and KIM-1 do not distinguish acute effects of calcineurin inhibitors in 2D monolayers or 3D perfused kidney MPS. (A) Primary PTECs in 2D were exposed to CsA or VCS for 48 hours across 0.01–10 µM (medium renewed at 24 h). Cell viability, assessed with an MTS tetrazolium assay, is normalized to matched vehicle control within donor and plotted as mean ± S.D. percentage across donors (n = 4). Two-way ANOVA with Šidák correction detected no significant main effect of drug or interactions between drug and concentration (adjust P > 0.05). (B) 3D perfused kidney MPS seeded with PTECs from the same donors were exposed to vehicle, 10 µM CsA, or 10 µM VCS for 48 h (medium refreshed at 24 h). KIM-1 in MPS effluents was quantified by ELISA at 0, 24, and 48 h and is shown as pg/mL, mean ± S.D. (n = 4). Two-way ANOVA with Šidák correction detected no significant main effect of treatment or interactions between treatment and time (adjust P > 0.05).

**Figure 3.**
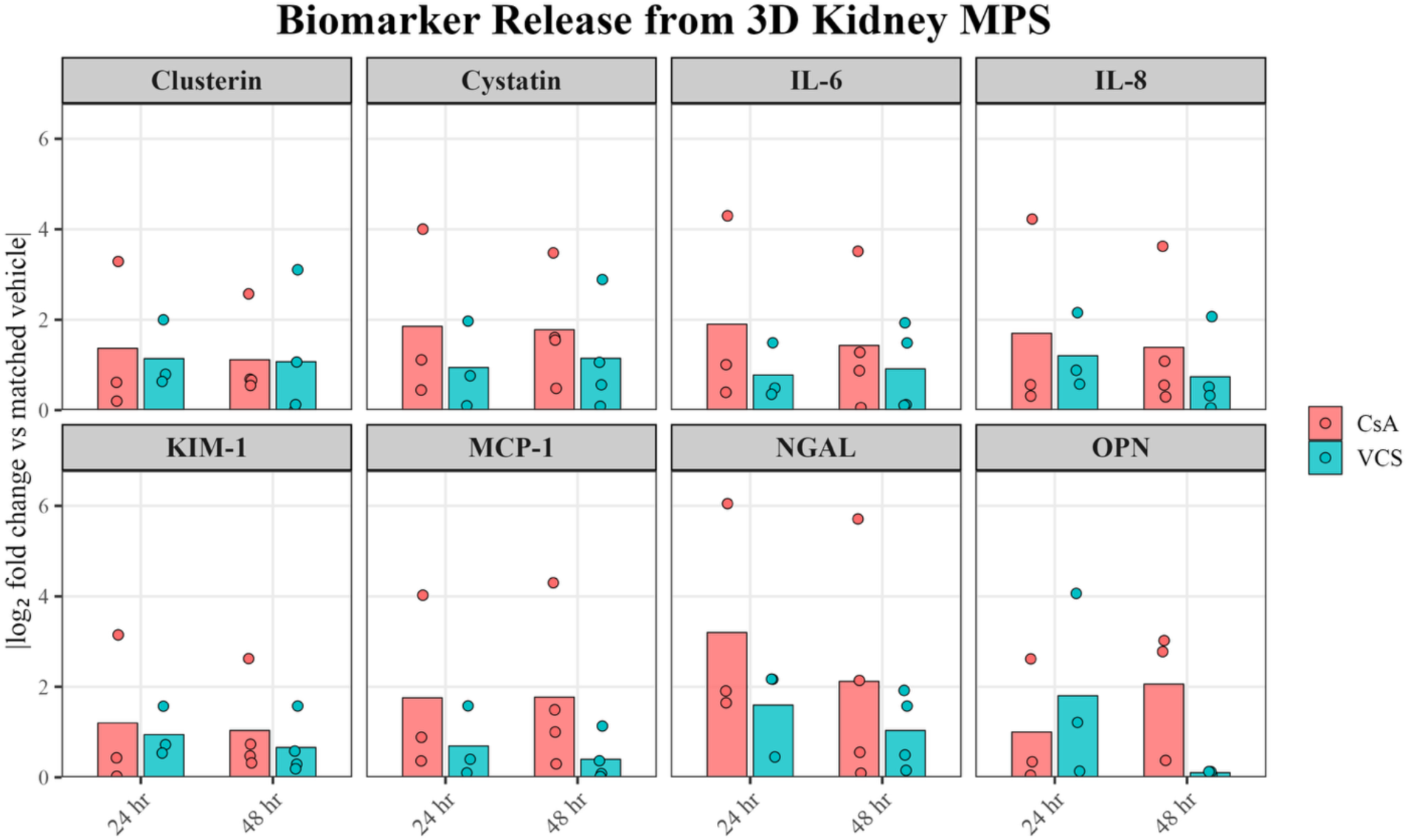
CsA produces greater aggregated MSD biomarker deviations than VCS in kidney MPS effluents. Bars show mean |log₂FC from matched vehicle| and points show individual donor values for eight injury and inflammatory biomarkers measured in MPS effluents after 24 and 48 h exposure to 10 µM CsA or 10 µM VCS. Larger values indicate greater absolute deviation from matched vehicle; 1 log₂ unit corresponds to a two-fold difference. The 24 h endpoint includes n = 3 donors for all biomarkers. The 48 h endpoint includes n = 4 donors for most biomarkers and n = 3 for OPN. Missing or non-computable matched values were omitted. Across biomarkers, after averaging 24 and 48 h endpoints within each biomarker, CsA produced greater absolute deviations than VCS (mean paired difference = 0.72 log₂ units; 95% CI = 0.37–1.07; paired t test, P = 0.0017; Wilcoxon signed-rank P = 0.0078). Data summarize the inter-donor means used in the mixed-effects analysis (Table S2, S3).

### CsA, but not VCS, disrupts mitochondrial architecture and membrane potential in perfused human proximal tubule MPS

To directly compare the two CNIs, we imaged 2D PTEC monolayers and 3D kidney MPS tubules after 48 h of exposure. In 2D PTEC monolayers, high-resolution confocal microscopy showed that vehicle-treated PTECs retained a continuous reticular network (Figure 4A). After 48 h of CNI treatment, CsA-treated cells exhibited a punctate pattern consistent with mitochondrial fission, while VCS-treated cells preserved filamentous morphology similar to control. Segmentation of individual mitochondria across 30 to 60 cells per donor (n = 4) showed no significant difference from vehicle in branch number or projected footprint for any group (Figure 4B, 4C). This lack of significant differences was also observed in a representative donor under hypoxic conditions across donors (Figures S2, S3). We then evaluated the same endpoints volumetrically in 3D kidney MPS tubules. Spinning-disk confocal tomography enabled full-volume reconstruction of perfused MPS tubules (Figure 4D). Filament area distributions differed modestly by treatment (Figure 4E). Turkey post-doc testing showed no significant difference between CNT and CsA, whereas VCS exceeded CNT (One-way ANOVA, Tukey’s correction, P *=* 0.0339). Mitochondrial membrane potential, quantified by the percentage of TMRM-positive cells, showed a treatment dependent change (Figures 4F, 4G). Across donors, both CsA and VCS exceeded vehicle (all pairwise comparisons vs CNT < 0.0001 shown above bars). VCS was greater than CsA in donors 2 and 3 (P = 0.0006 and P = 0.0023, respectively). Together, qualitative imaging shows CsA-associated fragmentation in 2D and 3D views, while group-level 3D filament area is not reduced versus vehicle. VCS maintains or even increases filament area and yields the highest TMRM-positive percentage. VCS preserved reticular mitochondrial architecture and yielded the highest TMRM-positive fraction, supporting preservation of mitochondrial membrane potential relative to CsA.

**Figure 4.**
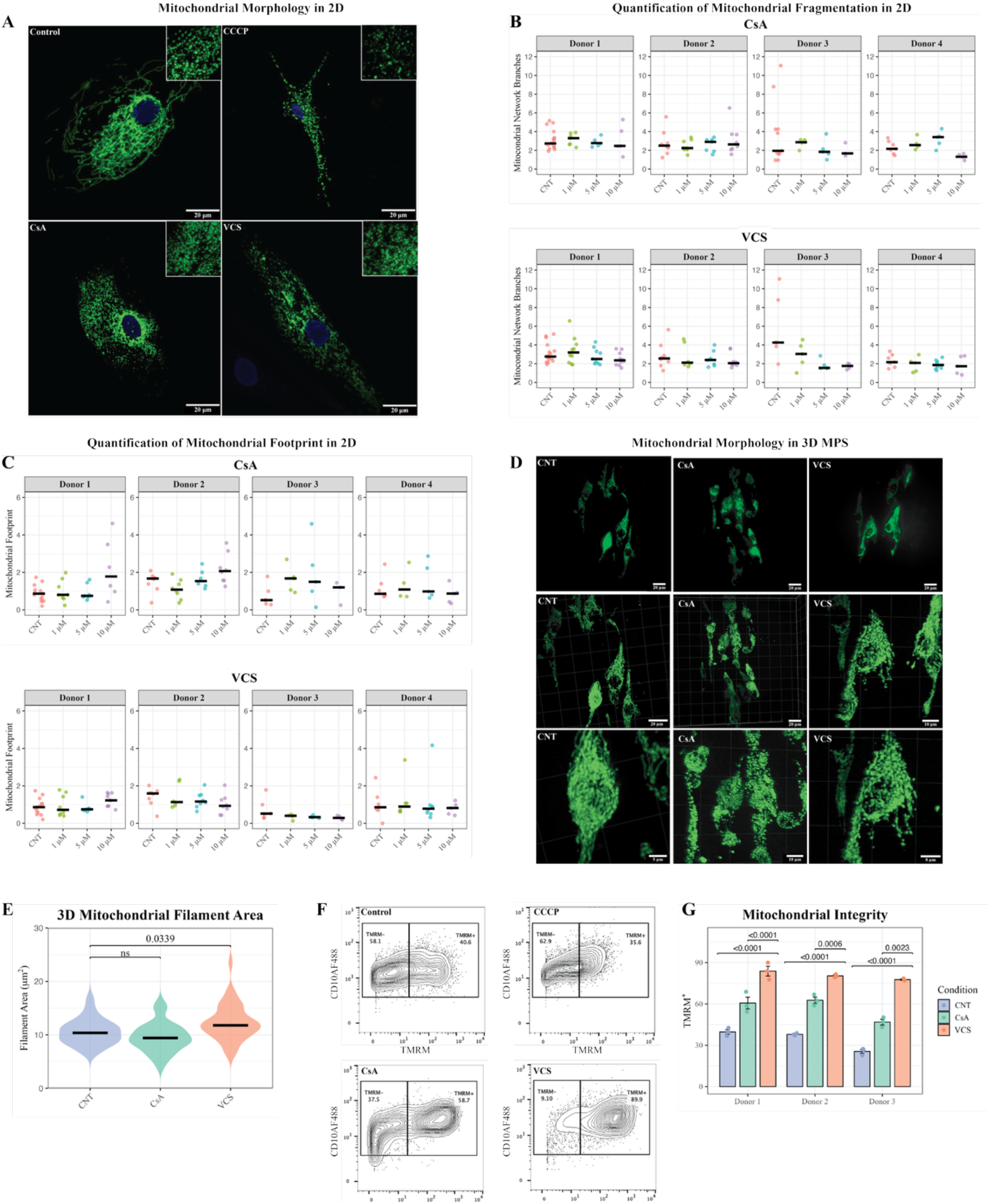
CNI effects on mitochondrial architecture and membrane potential in human PTECs. (A) Representative confocal images of 2D PTEC monolayers after 48 h exposure of vehicle (CNT), 10 µM CCCP (positive fragmentation control), 10 µM CsA, or 10 µM VCS. Mitochondria after CNT and VCS exposures show reticular networks, whereas that after CsA exposure appears punctate and fragmented. (B & C) Morphometrics of 2D PTECs (30 to 60 cells per donor, n = 4) after exposure to 1, 5 or 10 µM CsA or VCS. No significant differences of (B) mitochondrial network branches and (C) projected footprint versus CNT for either CNI were observed. (D) Spinning-disk confocal tomography of perfused 3D MPS tubules illustrates volumetric morphology under CNT, CsA, and VCS. (E) Violin plots summarizing 3D filament area from 105 reconstructed cells. One-way ANOVA detected a treatment effect (P = 0.0003); Tukey post-hoc testing showed that VCS had greater filament area than CsA (P = 0.0002), whereas CNT and CsA did not differ significantly. (F) Representative contour plots show tetramethylrhodamine methyl ester (TMRM) fluorescence in 2D PTEC cultures following 48 h exposure with CNT, 10 µM CCCP, 10 µM CsA, or 10 µM VCS. Numbers denote the percentage of TMRM-positive cells. (G) Percentage of TMRM-positive cells by donor (mean ± S.D., n = 3). Two-way ANOVA confirmed a treatment effect. TMRM-positive fraction was highest after VCS exposure; VCS exceeded CsA in donor-level comparisons for donors 2 and 3 (P = 0.0006 and P = 0.0023, respectively).

### CsA triggers a broad transcriptional remodeling, whereas VCS induces a constrained ER-chaperone response

Bulk RNA-seq of matched kidney MPS tubules harvested at 48 h showed markedly different transcriptome breadth between the two CNIs (Tables S4 to S6 and Figures S4 to S8). Using an FDR < 0.05 threshold, differential-expression analysis identified 1492 transcripts under CsA and 489 under VCS (Figure 5A). Comparison analysis revealed 1188 CsA-unique genes under CsA, 185 VCS-unique genes, and 304 shared genes (Figure 5B). Plotting the log₂-fold change for each gene under CsA against its VCS counterpart showed that the shared genes clustered tightly along the diagonal, whereas the CsA-exclusive genes deviated from this line, demonstrating the wider expression range of the CsA response (Figure 5C). KEGG analysis demonstrated that CsA preferentially enriched nuclear and cell-cycle modules, including DNA replication, mismatch repair, base-excision repair, and cell cycle (Figure 5D). Under VCS, enrichment was dominated by protein processing in the endoplasmic reticulum and related unfolded-protein-response branches (Figure 5D). Gene ontology analysis was consistent with these trends (Figures 5E, 5F). CsA upregulated biological-process terms linked to chromosome organization, chromosome segregation, mitotic cell cycle, and cell division, with cellular-component terms mapping to centromeric chromatin and kinetochore structure. VCS mainly increased ER quality control programs, including ER-chaperone complex, unfolded-protein binding, and ATP-dependent protein-folding, with more limited enrichment of nucleus- and cell-cycle-associated categories. Together, these transcriptomic data indicate that CsA engages a broad replication/repair and checkpoint-associated response, whereas VCS produces a narrower, primarily ER protein quality control signature.

**Figure 5.**
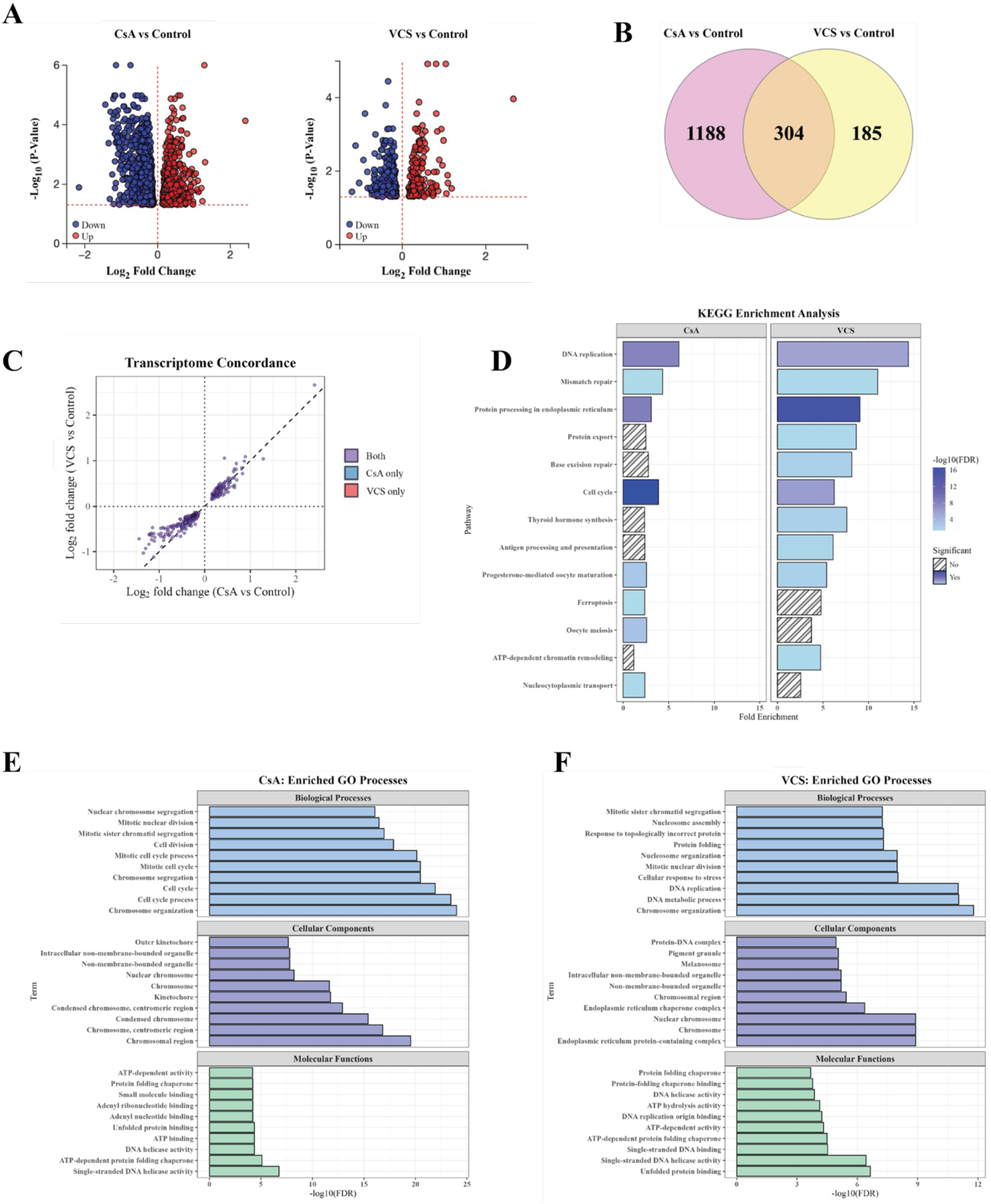
Global transcriptomic responses of PTECs to CNI exposure. (A) Volcano plots showing the log₂-fold change (x-axis) versus –log₁₀(adjusted P) (y-axis). Points passing the false-discovery threshold (FDR < 0.05) are colored red (up) or blue (down). (B) Venn diagram showing differentially expressed transcripts unique to CsA, unique to VCS, or shared by both. (C) Transcriptome concordance. For each transcript, log₂-fold change under CsA (x-axis) is plotted against VCS (y-axis). Dashed diagonal indicates identical. Purple points are significant for both drugs. Blue/red denote transcripts significant only with CsA or only with VCS, respectively. (D) KEGG pathway enrichment among regulated transcripts by CsA (left panel) or VCS (right panel). Bar length indicates fold enrichment. Color shading indicates –log₁₀(adjusted P). Hatched bars did not reach significance. (E & F) Gene Ontology enrichment for CsA (E) and VCS (F), organized by Biological Process, Cellular Component, and Molecular Function. X-axis shows – log₁₀(adjusted P). See Tables S4-S6 and Figures S4-S8 for a more detailed description of pathway analysis.

As an exploratory regulatory analysis, miRNA-target enrichment identified 88 miRNA target sets associated with the CsA differential-expression profile, whereas no VCS-associated miRNA target sets passed FDR correction (Figure S8). Because this analysis was based on differentially expressed mRNA targets rather than direct small-RNA measurement, these results are considered hypothesis-generating.

### CsA perturbs ER stress, cell-cycle, and ferroptosis pathways, whereas VCS mainly modulates ER protein-quality control

To resolve pathway-level differences, we visualized representative genes from ER quality control, cell-cycle regulation, and ferroptosis (Figure 6). Within the ER pathway, CsA increased luminal chaperones (*HSPA5/GRP78, HYOU1*) and induced UPR transcriptions factors *ATF6B* and *XBP1*. In contrast, several ER-associated degradation (ERAD) co-chaperones (*HSPA4L, HSPA8, HSP90AA1/AB1, HSPH1, DNAJB1/A1*) were modestly reduced, indicating a mixed ER-stress remodeling rather than a uniform upshift of ERAD. VCS, by comparison, preferentially increased ER chaperones and the ERAD lectin *EDEM1*, and upregulated *CRYAB*, without induction of upstream UPR transcription factors, which is consistent with a targeted, adaptive protein quality control response (Figure 6A). In the cell-cycle module, CsA upregulated *CDKN1A* (p21) and downregulated G2/M and spindle assembly regulators (*PLK1, CCNB1, TTK, CDC20, MAD2L1*), compatible with p21-mediated G2/M arrest (Figure 6B). VCS produced a milder, predominantly suppressive pattern across replication/checkpoint genes (*AURKB, CCNB1/2, PCNA, MCM2, TRIP13, MAD2L2*) without p21 induction, suggesting cell cycle deceleration rather than stable arrest (Figure 5B). Ferroptosis-related transcripts further distinguished VCS from CsA (Figure 5C). CsA increased lipid-metabolism genes (*SAT2, ACSL1*) and modestly altered iron/oxidative-damage-related nodes (increased *PCBP2*, decreased *TFRC*, *VDAC3*, *ATG7*), a pattern that is consistent with increased susceptibility to lipid peroxidation. VCS markedly induced ceruloplasmin (*CP*) and decreased *TFRC* and *VDAC3*, changes predicted to limit labile Fe²⁺ and restrain peroxidative stress. These data collectively indicate that CsA activates ER stress with cell-cycle arrest and a transcriptional pattern consistent with ferroptosis-priming, while VCS mainly refines ER quality control and exerts limited effects on cell-cycle or ferroptosis-related pathways in PTECs.

**Figure 6.**
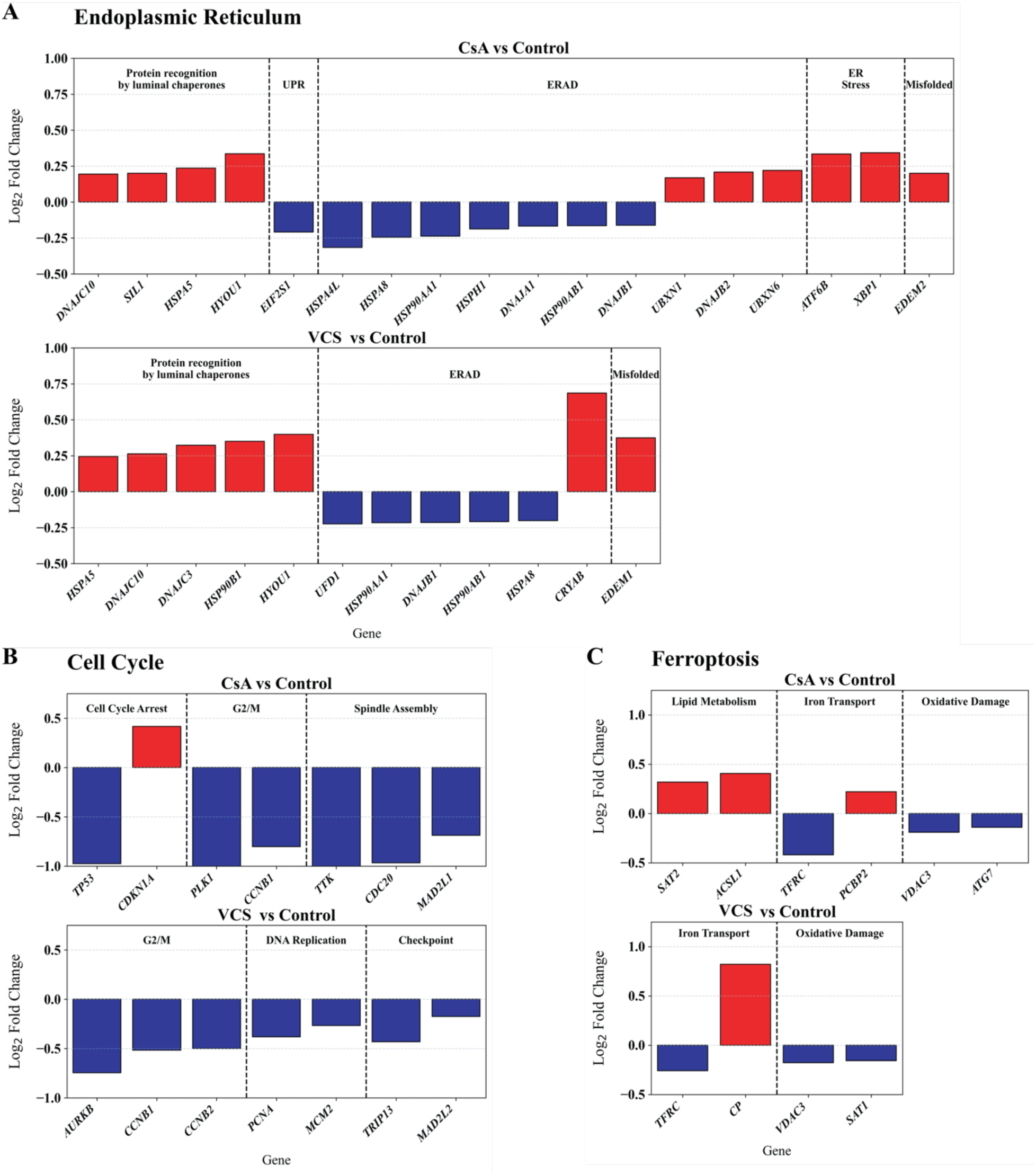
Targeted pathway analysis reveals distinct transcriptional signatures in PTECs after CNI exposure. (A) Representative ER quality-control genes for CsA (upper panel) and VCS (lower panel), grouped by functional sub-modules (dashed dividers): luminal chaperone recognition, UPR, ERAD, ER stress, and misfolded-protein handling. (B) Selected genes displayed as log₂-fold change versus control. CsA (upper panel) is arranged by cell-cycle arrest, G2/M transition, and spindle assembly. VCS (lower panel) is arranged by G2/M, DNA replication machinery, and checkpoint regulation. (C) Key genes governing lipid-peroxidation metabolism, iron transport, and oxidative-damage response for CsA (upper panel) and VCS (lower panel). Functional groupings are indicated above each sub-panel. Red = upregulated, blue = downregulated. Only differentially expressed genes are shown (FDR < 0.05). Horizontal baselines indicate no change (log₂-fold change = 0).

## Discussion

CNIs remain indispensable for rejection prophylaxis after solid-organ transplantation, but the nephrotoxicity associated with first-generation agents such as CsA continues to limit long-term allograft survival. VCS, a single–amino-acid analog of CsA, has shown a more favorable renal safety profile in clinical studies, yet the cellular basis for this distinction has not been fully defined (Figure S9). Here, using a perfused, 3D human kidney MPS coupled with multimodal readouts, we identified fundamental differences in how CsA and VCS affect PTECs. Notably, these differences are not apparent in static monolayers, highlighting the importance of physiologically relevant culture systems for uncovering sub-lethal drug toxicities.

Initial benchmarking in static PTEC monolayers revealed no overt toxicity. Neither agent reduced tetrazolium-based cell viability nor increased KIM-1 over 48 hours. However, the same cells subjected to physiologic flow within a 3D MPS revealed a hidden vulnerability to CsA. Confocal imaging demonstrated CsA-associated mitochondrial network fragmentation, whereas VCS preserved reticular architecture and yielded a higher TMRM-positive fraction than CsA. These observations are consistent with mitochondrial permeability transition events and ROS overload, phenomena linked to tubular stress^11–13,18–21^. In contrast, VCS preserved a reticular mitochondrial network structure and sustained ΔΨm at matched molar exposure of CsA. Together, this pattern is consistent with ROS-triggered, cyclophilin-D-dependent mitochondrial permeability transition pore engagement by CsA^22–24^, whereas VCS’s single amino-acid substitution may attenuate this mitochondrial liability while retaining calcineurin-inhibition potency.

Transcriptomic profiling linked these organellar events to proteostasis disruption (Figure 6A). In CsA-treated PTECs cultured in 3D MPS, canonical UPR sensors (*ATF6B, XBP1*) were induced together with luminal chaperones (*HSPA5/GRP78, HYOU1*), while multiple ERAD co-chaperones (*HSPA4L, HSPA8, HSP90AA1, HSPH1, DNAJ* family) and *EIF2S1* were reduced, with only a modest rise in *EDEM2*. This pattern suggests a mixed, partly maladaptive UPR, with heightened chaperone demand with blunted translation attenuation and reduced ERAD-associated genes^25–27^. In contrast, VCS increased ER-chaperone and the ERAD lectin *EDEM1* and upregulated CRYAB without inducing *ATF6B/XBP1*, consistent with a targeted, adaptive protein quality control response^28,29^.

Concomitant analysis further highlighted CsA’s propensity to prime ferroptosis (Figure 6C). CsA increased lipid-metabolism nodes (*SAT2*, *ACSL1*) and shifted iron/oxidative-damage modules (increased *PCBP2*, decreased *TFRC*, *VDAC3*, and *ATG7*) toward a state permissive for lipid peroxidation. VCS, by comparison, suppressed TFRC and strongly induced CP, while reducing VDAC3, which are changes predicted to limit labile Fe²⁺ and restrain peroxidative stress^30–32^.

A third axis involved a broad transcriptional blockade of cell-cycle progression (Figure 6B). In PTECs treated with CsA, *CDKN1A* (p21) was induced and G_2_/M and spindle-assembly regulators (*PLK1*, *CCNB1*, *TTK*, *CDC20*, *MAD2L1*) were downregulated, which are compatible with p21-mediated G_2_/M arrest^33^. VCS produced a milder, predominantly suppressive pattern across replication/checkpoint genes (*AURKB*, *CCNB1/2*, *PCNA*, *MCM2*, *TRIP13*, *MAD2L2*) without p21 induction, implying slowed cell cycling rather than stable arrest. Analysis from these three axes collectively show that CsA activates ER stress with checkpoint-enforced arrest and a ferroptosis-prone shift, whereas VCS mainly increases ER protein quality control and diminishes ferroptosis priming at matched exposure.

An exploratory miRNA-target enrichment analysis provided a hypothesis-generating overlay to the RNA-seq findings. Target sets for 88 miRNA regulators were enriched in the CsA differential-expression profile, whereas no VCS-associated miRNA target sets passed FDR correction (Figure S8). Because this analysis inferred miRNA-regulatory associations from mRNA target enrichment rather than directly measuring miRNA abundance, these results should be interpreted as predicted regulatory nodes. Future studies using dedicated small-RNA sequencing or targeted miRNA assays in cell lysates, effluents, or plasma should determine whether these computationally nominated miRNA regulators are measurable biomarkers of early CNI-associated tubular stress.

Another important observation is the discordance between organellar and transcriptional stress and the absence of a KIM-1 signal at 48 hours. This highlights a biomarker gap. KIM-1 is valuable for detecting overt proximal-tubule injury, yet our data show it may not report sub-lethal stress states that nevertheless reduce repair capacity or prime ferroptosis^34^. A panel that integrates mitochondrial readouts with ER-stress, ferroptosis transcripts, and repaired-capacity transcripts could provide earlier and mechanistically based signals for dose adjustment or CNI selection. Exploratory miRNA-target enrichment may provide additional regulatory hypotheses, but direct small-RNA profiling will be needed before nominating miRNAs as biomarkers. In a clinical setting, such a panel might enable proactive switching from CsA to VCS or dose adjustments before irreversible renal tubular injury occurs.

Our findings also emphasize the importance of physiologic culture context for revealing nuanced toxicology. The 3D MPS introduces shear stress and flow-driven nutrient and drug gradients that are absent in static monolayers. These features allow us to stress-test the energy intensive liabilities of protein folding and trafficking that are only evident under flow. Of note, the four primary PTEC donors exhibited the natural variability seen clinically (two-to three-fold differences in baseline viability, IL-6 secretion, and the absolute decrease in TMRM), yet the direction and relative magnitude of drug effects (CsA > VCS ≈ vehicle) were conserved across donors. This reproducibility despite biological heterogeneity suggests that the mechanistic separation we observe is robust and supports further testing across broader donor cohorts.

Our study has a few limitations that outline clear directions for future work. First, we examined PTECs in isolation. Glomerular and vascular compartments, which contribute to hemodynamic components of CNI toxicity, were not included. Multicellular kidney-chip platforms that incorporate peritubular capillary endothelium and circulating immune cells could delineate how endothelial-tubular crosstalk and immune-cell interactions could affect the pathways we describe^35,36^. Second, our 48-hour window captures early events. Longer perfusions with repeated sampling will be necessary to determine whether transient ΔΨm loss and ferroptosis priming predict sustained tubular injury or interstitial scarring. Third, we inferred pathway activation from bulk RNA-seq and miRNA-regulatory associations primarily from bulk RNA-seq. Single-cell transcriptomics, proteomics, and direct assays of lipid peroxidation and iron redox state, and dedicated small-RNA profiling will provide further validation of our findings.

In conclusion, despite both agents inhibiting calcineurin, CsA and VCS differentially shape the tubular stress landscape in human PTECs under physiologic flow. CsA disrupts mitochondrial architecture, propagates maladaptive ER stress, primes ferroptosis, and enforces p21-mediated arrest, whereas VCS induces adaptive ER quality-control pathways while preserving mitochondrial integrity and proliferative capacity. These mechanistic distinctions align with the clinical renal-sparing profile of VCS and provide a biologic rationale for agent selection in patients at high risk for nephrotoxicity. Notably, our work also illustrates how perfused human kidney-chip models, paired with multimodal readouts, can reveal sub-lethal nephrotoxic pathways before injury markers rise in conventional systems and assays.

## Funding

This work was supported in part by the National Institutes of Health (NIH) National Center for Advancing Translational Sciences (NCATS) awards UG3TR002158 and TL1TR002318, the National Institute of Environmental Health Sciences (NIEHS) Interdisciplinary Center for Exposures, Diseases, Genes and Environment (UW EDGE Center, P30ES007033), and the University of Washington School of Pharmacy, Department of Pharmaceutics Ji-Ping Wang award. Aurinia Pharmaceuticals Inc. provided research funding support for these studies.

## Acknowledgments

We thank Dr. Dale W Hailey at the UW Institute for Stem Cell and Regenerative Medicine for his assistance with mitochondrial imaging. We also thank Gohar Mosoyan at MSSM for her assistance with the MSD biomarker/cytokine assays.

## Author Contributions

Conceptualization: **Aryeh, Yeung, Himmelfarb, Kelly**

Data curation: **Aryeh, Tsang, Hsu, MacDonald, Bammler, Rehaume**

Formal analysis: **Aryeh, Tsang, Hsu, MacDonald, Bammler, Rehaume**

Funding acquisition: **Yeung, Himmelfarb, Rehaume, Kelly**

Investigation: **Aryeh, Tsang**

Methodology: **Aryeh, Tsang, Hsu, Yeung, Kelly, Himmelfarb**

Supervision: **Yeung, Kelly, Himmelfarb**

Writing – original draft: **Aryeh, Tsang**

Writing – review and editing: **Hsu, Yeung, MacDonald, Bammler, Himmelfarb, Rehaume, Kelly**

## Data Sh**a**ring Statement

RNA-sequencing data have been deposited in the National Center for Biotechnology Information (NCBI) Gene Expression Omnibus (GEO) under accession number GSE309956.

## Notes

### Competing Interest Statement

Aurinia Pharmaceuticals Inc. provided research funding support for these studies. Linda M. Rehaume was affiliated with Aurinia Pharmaceuticals Inc. during the conduct of the work. The remaining authors declare no competing interests.

