## Supplementary Material for "Voclosporin Preserves Mitochondrial Function Compared With Cyclosporine A in Perfused Human Proximal Tubule Microphysiological Systems"

### SUPPLEMENTAL MATERIAL:

This article contains the following supplemental material:

**1-12. Supplemental Methods**

13. Summary of Demographic and Clinical Characteristics of Human Donors.................... 10

14. KIM-1 expression showed no significant change in 3D PTECs treated with CsA and VCS at 48 h and 14 d ........ 11

15. Linear mixed-effects model analysis of biomarker responses following treatment ........ 12

16. Paired t-test comparison of $log_2$ biomarker responses ........ 14

17. Mitochondrial Morphology of 2D-cultured PTECs Under Normoxic and Hypoxic Conditions Following CsA and VCS Treatments ........ 15

18. Mitochondrial footprint and network branching in 2D-cultured PTECs under normoxic and hypoxic conditions following CsA or VCS treatment ........ 16

19. Summary of RNA-seq Results for CsA and VCS ........ 17

20. Pathway enrichment analysis for CsA treatments ........ 18

21. Pathway enrichment analysis for VCS treatments ........ 19

22. Cell cycle pathway analysis in PTECs treated with CsA ........ 20

23. Cell cycle pathway analysis in PTECs treated with VCS ........ 21

24. ER pathway analysis in PTECs treated with CsA ........ 22

25. ER pathway analysis in PTECs treated with VCS ........ 23

26. Differential miRNA expression in CsA and VCS treatments compared to control ........ 24

27. Mechanism of calcineurin inhibition by CsA and VCS ........ 25

**Supplemental Methods**

### 1) Chemicals and Reagents

Dulbecco's Modified Eagle's Medium/Nutrient Mixture F-12 powder without glucose was purchased from United States Biological (Salem, MA, USA). D-Glucose, and HEPES (4-(2-hydroxyethyl)-1-piperazineethanesulfonic acid), sodium bicarbonate. Dulbecco's Phosphate-Buffered Saline (DPBS), Insulin-Transferrin-Selenium (100X) (100X ITS-G), Antibiotic–Antimycotic (100X) (100X AA), Trypsin-EDTA, Defined Trypsin Inhibitor (DTI), Collagenase type IV, and Dispase® powder was purchased from Gibco (Billings, MT, USA). 2-[2-(3-chlorophenyl)hydrazinylidene]-propanedinitrile (CCCP) was purchased from Cayman (Ann Arbor, MI, USA). Polybrene was purchased from Millipore Sigma (USA). **Cyclosporin A (CAS 59865-13-3**, Lot A0422) and **Voclosporin (CAS # 515814-01-4)** were purchased from Chem Cruz (Dallas, TX, USA).

### 2) Primary Human PTEC Isolation and Culture

Human PTECs were isolated from normal human kidneys (GFR > 60 mL/min) obtained through U.S. Organ Procurement Organizations via Novabiosis, Inc. (Durham, NC, USA). The kidneys were identified as non-transplantable and sourced from adult donors (S13). Kidneys were maintained at cold temperatures with clamp-to-isolation time of less than 36 hours. Under sterile conditions, the renal cortex was dissected and minced into approximately 1 $mm^3$ pieces. Minced tissue was enzymatically digested in sterile-filtered DPBS containing 0.75 mg/mL collagenase type IV and 0.75 mg/mL dispase, with gentle agitation for 1 hour at 37°C on an orbital shaker. The resulting cell suspension was passed through a 100 μm cell strainer, and the filtrate was centrifuged at 200 x g for 7 minutes. All subsequent cell handling steps were performed at 4°C. Cell pellets were washed twice with cold PBS and resuspended in warm PTEC media before plating in a 75 $cm^2$ flask. PTECs were expanded to confluency and cryopreserved for use in 2D (Lentivirus biosensor transduction, MTS) and 3D culture systems (MPS).

#### 3) PTEC Seeding in 3D MPS

Detailed protocol for PTEC seeding in 3D MPS platform has been previously described and validated (Weber et al., 2016). In brief, the 3D MPS platform, or "chip," was developed by Nortis Inc. (Bothell, WA, USA). For PTEC seeding, chips were filled with 6 mg/mL rat tail collagen type I (Ibidi, Madison, WI) at 4°C. The chips were incubated for 30 minutes at 4°C to allow the matrix to settle and subsequently stored at room temperature overnight. Microfiber inserts were removed from the device, and the channel was coated with 5 µg/mL mouse collagen type IV (Corning, NY, USA) at 37°C for 1 hour.

Cryopreserved PTECs were thawed and cultured in 75 $cm^2$ flasks until confluency. Confluent monolayers of PTECs were trypsinized with Trypsin-EDTA to obtain single-cell suspensions. PTEC pellets were then washed with DTI and resuspended at a density of 15 to 20 million cells/ml. Approximately 5 µL of cell suspension was injected into one collagen IV-coated lumen of the MPS. Cells were allowed to adhere for 24 hours before initiating 0.5 µL/min of media flow. Cell coverage and integrity of the tubule structure were assessed under light microscopy weekly. For all experiments conducted in this study, PTECs in chips were grown for 1 to 2 weeks after seeding.

#### 4) CsA and VCS Treatments

To mimic acute drug exposures, PTECs in both 2D and 3D culture systems were treated with CsA and VCS at 0, 1, 5, and 10 µM for 48 hours. Media was replenished every 24 hours. Human serum albumin (HSA) was included in the treatments at 100 ug/mL. The percentage of organic solvents were normalized across all treatment groups to maintain consistency. For chronic drug exposure, CsA and VCS were dosed for 14 days with media changes performed at 24 hours, 48 hours, and day 7 or at day 4 and 10.

#### 5) Cell Viability Assessment via MTS Assay

To evaluate the effects of CsA and VCS on PTEC viability, a colorimetric MTS assay was performed. PTECs were seeded in 96-well plates and allowed to adhere overnight under standard culture conditions. Cells were then treated with CsA or VCS at final concentrations ranging from 0.01 to 10 µM for 48 hours, with drug-containing media replenished at 24 hours. Following the 48-hour incubation, MTS reagent (3-(4,5-dimethylthiazol-2-yl)-5-(3-carboxymethonyphenyl)-2-(4-sulfophenyl)-2H-tetrazolium; ab197010, Abcam) was added to each well according to the manufacturer's protocol and incubated for 3 hours at 37°C. Absorbance was measured at 490 nm using a Multiskan SkyHigh Microplate Spectrophotometer (Thermo Fisher Scientific). Viability was expressed as a percentage relative to untreated controls. Each condition was tested in technical triplicates, and the experiment was repeated using cells from multiple donors (n = 4). Data are presented as mean ± SD.

#### 6) Mitochondrial Labeling Using Lenti Virus Biosensor Transduction

Mitochondrial labeling was conducted in both 2D and 3D cultured PTECs to assess mitochondrial integrity following drug treatments. PTECs at ~50% confluence were exposed to 10 µg/mL

Polybrene (MilliporeSigma, St. Louis, MO) and transduced with a pCT-COX8-GFP mitochondrial targeting construct (multiplicity of infection [MOI] = 10:1) provided by Dr. Lansing Taylor of Pittsburg.

After 72 hours, the culture medium was refreshed. Successful mitochondrial labeling was confirmed via fluorescence microscopy (Nikon Ti-E). PTECs were fixed in 4% paraformaldehyde (PFA), and their mitochondrial network morphologies was quantified using ImageJ/Fiji with the MiNA plugin or Imaris.

#### 7) PTEC Genotyping

Genotyping was performed using TaqMan™ SNP assays (Applied Biosystems, Foster City, CA, USA) to verify the genetic identity and stability of the PTECs. Each DNA sample was mixed with TaqMan™ Genotyping Master Mix (Applied Biosystems) and specific TaqMan™ SNP assay probes in 96-well plates. The PCR amplification and allelic discrimination were carried out using a QuantStudio Real-Time PCR System (Applied Biosystems) under standard cycling conditions according to the manufacturer's guidelines. Post-run analysis was conducted with QuantStudio Design and Analysis Software to ensure the presence of expected SNP alleles and to confirm the consistency of the genetic profiles across cell passages.

#### 8) KIM-1 Measurements of 3D PTEC Effluents

To assess the levels of kidney injury molecule-1 (KIM-1; HAVCR1) in 3D PTEC cultured effluents, samples were collected at predetermined intervals (for short-term studies: 0, 24, and 48 hours; for long-term studies: 4, 7, 10, and 14 days). KIM-1 protein concentrations were measured using a DuoSet ELISA kit (R&D Systems), following the manufacturer's instructions. Briefly, undiluted effluents were added to ELISA plates pre-coated with the KIM-1 capture antibody, then incubated with a detection antibody and streptavidin-HRP. After washing, substrate solution was added, and the reaction was stopped with the recommended stop solution. Absorbance was recorded at 450 nm using a microplate reader (Multiskan SkyHigh Microplate Spectrophotometer, Thermo Fisher Scientific), and KIM-1 levels were calculated against a standard curve generated from known concentrations of recombinant KIM-1.

**9) Spinning Disk and Confocal Microscopy**

Fluorescence imaging was conducted for both 2D and 3D samples using a Nikon A1R confocal microscope equipped with a 40x water objective to acquire Z-stacks for 3D reconstructions. Live-cell imaging in a spinning-disk configuration was performed for additional observations to capture dynamic changes in mitochondrial morphology in 3D. All images were processed in ImageJ/Fiji, and mitochondrial networks were quantified using the MiNA plugin.

Deconvolution of the images was performed using NIS-Elements software, and Z-stack images were reconstructed to visualize 3D tubule structures. Imaris software was utilized for advanced 3D visualization and analysis.

**10) Flow Cytometry**

Cells were seeded in 12-well plates, grown to ~80% confluence, and treated for 24 hours with vehicle, CCCP, or 10 µM drug CsA or VCS. PTECs were harvested by trypsinization, washed, then resuspended in 100 µL of 1× DPBS containing a fixable viability dye (1:1000 dilution), anti-

CD10 (MA5-27893, final 2 μg/mL), TMRM (ThermoFisher M20036, final 40 nM) and incubated at 37 °C for 20 minutes. CCCP (1 μM) was used as the positive control. Cells were washed, resuspended in 100 μL of 1× DPBS containing Goat-anti-rabbit Alexa Fluor™ 488 (ab150077, final ~2 μg/mL), incubated for 10 minutes at 37 °C, then washed again and analyzed on a BD A3 Symphony flow cytometer. Flow cytometry data were processed using FlowJo™ X software (Tree Star).

**11) MSD Biomarker Analysis**

Biomarkers and cytokines were quantified using a proprietary, analytically validated multiplex format on the MesoScale platform (MesoScale Diagnostics, Rockville, MD, USA), which employs electrochemiluminescence detection and patterned arrays for multiplex analysis. Each chip effluent sample was analyzed in duplicate, with all coefficients of variations (CVs) falling within acceptable ranges. In the Proinflammatory Panel 1, IL-6 and IL-8 were successfully quantified, while other targets in the panel were below the detection limits. NGAL, cystatin C, and clusterin were detected in all samples. KIM-1 and MCP-1 were detected reliably, whereas EGF, IL-11, αGST, and IL-18 fell below the assay's lower limit of detection. The complete targets in the panel included: IFN-γ, IL-10, IL-12p70, IL-13, IL-1β, IL-2, IL-4, IL-6, TNF-α, EGF, IL-11, IL-18, KIM-1, MCP-1, OPN, αGST, clusterin, cystatin C, and NGAL.

**12) RNA-sequencing, Transcriptomic Analysis, and Meta-Analysis**

Total RNA was extracted from MPS-cultured PTECs (n = 3) after 48-hour drug treatments using a Qiagen RNeasy Micro Kit, and quality was verified using an Agilent Bioanalyzer 2100. Libraries were prepared with the SMARTer Stranded Total RNA Sample Prep Kit - Low Input Mammalian

(Takara Bio, San Jose, CA, USA) and sequenced on Illumina NovaSeq 6000 and NovaSeq Plus X platforms. Differential expression analysis was performed using the limma-voom pipeline, either fitting a linear mixed model with a random intercept for each subject or stratified by subject. Genes were selected using a Benjamini-Hochberb false discovery rate (FDR)<0.05. . Pathway enrichment analyses were conducted using KEGG and iPathwayGuide. iPathwayGuide employs an impact analysis approach, integrating gene-level changes, pathway topology, and the roles of each gene within the network to identify biologically meaningful pathway perturbations. Publicly available KEGG pathway datasets were included for meta-analysis and contextualization of results. Additionally, differentially expressed genes were further examined with ENRICHR and BioJupies to identify key pathways, regulatory networks, and upstream regulators. Exploratory miRNA-target enrichment analysis was performed in iPathwayGuide. The analysis tested whether predicted or curated targets of known miRNAs were overrepresented among treatment-specific differentially expressed transcripts. Because no dedicated small-RNA sequencing or targeted miRNA assay was performed, these results are interpreted as predicted miRNA-regulatory enrichment rather than direct miRNA expression.

13. Table S1. Summary of Demographic and Clinical Characteristics of Human Donors

Summary of donor demographics, genotypes, and clinical histories.

| Donor ID | CYP3A5 Genotype | Sex | Age (years) | Ethnicity | Pre-existing Conditions | Cause of Death |
|---|---|---|---|---|---|---|
| **pT 2** | *3/*6 | F | 62 | White | – | – |
| **pT 3** | *3/*3 | F | 42 | White | History of opioid use | Opioid overdose |
| **pT 5** | NC | F | 47 | Hispanic | – | – |
| **pT 6** | *3/*3 | F | 45 | White | Hypertension, history of heavy alcohol use | Cerebrovascula r stroke |
| **pT 8** | *1/*7 | F | 51 | Black | – | – |
| **pT 9** | NC | M | 38 | White | – | – |
| **pT 10** | NC | M | 49 | White | Diabetes, hypertension, chronic | Cardiovascular anoxia |
| **pT 11** | *3/*3 | M | 64 | White | Diabetes, hypertension, coronary artery disease, history of cigarette use | Stroke, intracranial hemorrhage |
| **pT 13** | *3/*3 | M | 36 | White | – | – |
| **pT 16** | *3/*3 | M | 59 | White | – | Head trauma/blunt injury |
| **pT 17** | *3/*3 | F | 50 | - | – | – |
| **pT 19** | *3/*3 | F | 25 | White | – | – |
| **pT 20** | NC | F | 65 | Black | – | – |
| **pT 21** | *3/*3 | F | 52 | White | – | Cerebrovascula r stroke |
| **pT 23** | *3/*3 | M | 73 | White | Diabetes, hypertension | Natural causes, anoxia |
| **pT 24** | *3/*3 | - | - | - | – | – |
| **pT 25** | *3/*3 | F | - | - | – | – |

Summary of the sex, age, ethnicity, CYP3A5 genotype, relevant pre-existing medical conditions, and reported cause of death for each PTEC donor used in this study. Blue highlights indicate donors used for RNA sequencing. Missing or unreported data are denoted by "–". CYP3A5 genotypes were determined by TaqMan™ assay; three donors could not be classified (NC) due to inconclusive results.

14. Figure S1. (Top figure) KIM-1 expression showed no significant change in 3D PTECs treated with CsA and VCS at 48 h and 14 d. KIM-1 release into the effluent from 3D PTEC MPS systems treated with CsA or VCS was measured up to 14 days. No significant changes in KIM-1 release were detected across conditions (ANOVA, $p > 0.05$). (Bottom figure) Pooled data from all donors comparing CsA and VCS did not exhibit statistically significant differential impact on PTEC viability at any tested concentrations (two-way ANOVA with Šidák correction).

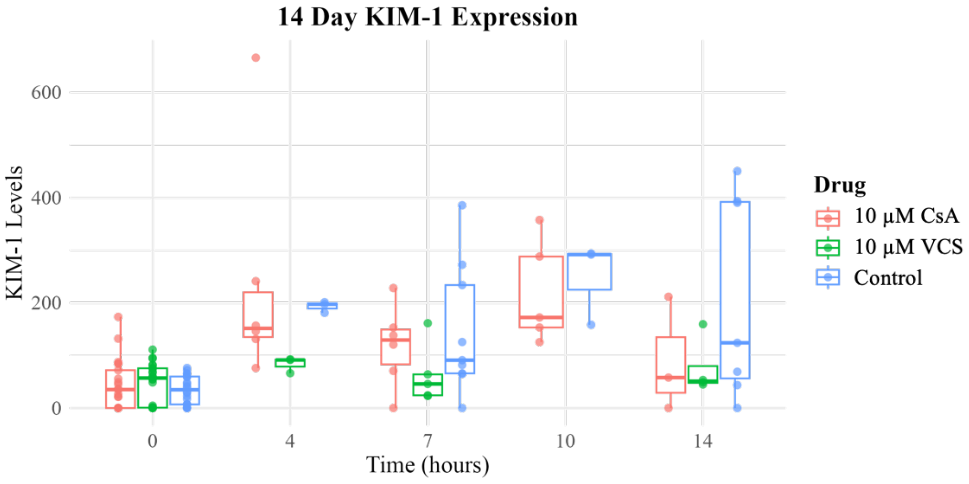


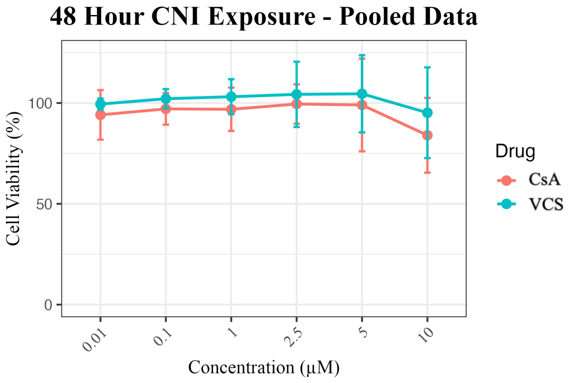

15. Table S2. **Donor level MSD biomarker values used for Figure 3.** Matched effluent values are shown for each biomarker, donor, time point, and treatment. Fold change was calculated relative to the matched vehicle control from the same donor and time point, and $log_2FC$ was calculated as $log_2$ (drug/matched vehicle). Figure 3 plot $|log_2FC|$ values at 24 and 48 h. Missing or non-computable matched values were omitted from summary statistics. The only non-computable 24/48 h value was OPN at 48 for donor pT6 because the matched CNT value was missing.

| Biomarker | Donor | Time | CNT | CsA | VCS | log2FC_CsA | log2FC_VCS |
|---|---|---|---|---|---|---|---|
| **Clusterin** | pT10 | 24 h | 856.7994743 | 983.54147 | 3422.228661 | 0.1990282886126450 | 1.9979066587196400 |
| **Clusterin** | pT23 | 24 h | 3929.802515 | 6014.314704 | 6811.303188 | 0.6139435457816000 | 0.7934740372219470 |
| **Clusterin** | pT6 | 24 h | 7479.49066 | 766.7638183 | 4824.856987 | -3.2860858620019300 | -0.6324538482277450 |
| **Clusterin** | pT10 | 48 h | 632.260662 | 1015.998801 | 5433.795435 | 0.6843073337557490 | 3.1033688879304200 |
| **Clusterin** | pT19 | 48 h | 13933.8 | 20339.4 | 15177.3 | 0.5456883604864400 | 0.1233264013872500 |
| **Clusterin** | pT23 | 48 h | 4146.840026 | 6566.53188 | 8666.288877 | 0.6631192179463480 | 1.063401934362910 |
| **Clusterin** | pT6 | 48 h | 5512.469014 | 927.4830471 | 5527.792863 | -2.57130582425681 | 0.004004914992597140 |
| **Cystatin C** | pT10 | 24 h | 404.2077684 | 187.1257705 | 1580.475309 | -1.1110887922263700 | 1.9671895428867500 |
| **Cystatin C** | pT23 | 24 h | 477.8751598 | 649.7480494 | 809.5107439 | 0.4432466201154940 | 0.7604164507414750 |
| **Cystatin C** | pT6 | 24 h | 2616.257514 | 163.4510764 | 2442.36953 | -4.000573766911930 | -0.09922305366858050 |
| **Cystatin C** | pT10 | 48 h | 495.6159905 | 162.7706091 | 3665.55615 | -1.606382514287840 | 2.886737465013490 |
| **Cystatin C** | pT19 | 48 h | 1201.4 | 3511.2 | 1276.5 | 1.547247606333170 | 0.08747696876801120 |
| **Cystatin C** | pT23 | 48 h | 807.9008926 | 1127.214486 | 1192.066232 | 0.48051182754730400 | 0.5612141655125340 |
| **Cystatin C** | pT6 | 48 h | 1826.202465 | 164.0455315 | 3799.582612 | -3.476678519622040 | 1.056994224752940 |
| **IL-6** | pT10 | 24 h | 37.865868 | 18.86078729 | 106.3240912 | -1.005508101537240 | 1.4894986185236900 |
| **IL-6** | pT23 | 24 h | 60.18644315 | 79.04828736 | 84.39588056 | 0.3932956458325550 | 0.48773402136112200 |
| **IL-6** | pT6 | 24 h | 243.8900555 | 12.41139847 | 191.2238478 | -4.296493346909390 | -0.3509684790312540 |
| **IL-6** | pT10 | 48 h | 47.31362095 | 25.8511364 | 132.6320035 | -0.8720278737851120 | 1.4871014524014800 |
| **IL-6** | pT19 | 48 h | 93.5 | 226.8 | 356.5 | 1.2783823701672300 | 1.9308638065562500 |
| **IL-6** | pT23 | 48 h | 109.0837196 | 113.3124481 | 100.1958813 | 0.054870559120727800 | -0.12261259442061900 |
| **IL-6** | pT6 | 48 h | 169.1698394 | 14.8334787 | 182.1403425 | -3.51154350036238 | 0.10657812292337400 |
| **IL-8** | pT10 | 24 h | 73.44158723 | 91.21782219 | 327.1251005 | 0.3127184876894480 | 2.155173318569770 |
| **IL-8** | pT23 | 24 h | 189.504521 | 279.6543799 | 282.5185924 | 0.5614126586369440 | 0.5761135465427360 |
| **IL-8** | pT6 | 24 h | 1204.263868 | 64.45757478 | 654.3660821 | -4.223657819574950 | -0.8799816619307770 |
| **IL-8** | pT10 | 48 h | 93.13784661 | 136.6207995 | 390.6014942 | 0.5527377080511980 | 2.0682580362139000 |
| **IL-8** | pT19 | 48 h | 168.5 | 356.5 | 134.9 | 1.0811534852620300 | -0.3208582431209530 |
| **IL-8** | pT23 | 48 h | 244.0308337 | 299.4831784 | 252.2724336 | 0.2954115246792990 | 0.04791912204578160 |

| | | | | | | | |
|---|---|---|---|---|---|---|---|
| **IL-8** | pT6 | 48 h | 883.2682025 | 71.70692519 | 618.1811607 | -3.6226672147325300 | -0.5148218874595620 |
| **KIM-1** | pT10 | 24 h | 4.605539407 | 4.532525394 | 13.69917464 | -0.0230550337468581 | 1.5726469333181500 |
| **KIM-1** | pT23 | 24 h | 22 | 29.66666667 | 36.33333333 | 0.43133931177004500 | 0.7237902052861160 |
| **KIM-1** | pT6 | 24 h | 24.71298968 | 2.785941163 | 35.83397239 | -3.149032858049280 | 0.536058430196874 |
| **KIM-1** | pT10 | 48 h | 8.58083074 | 6.182008062 | 25.59187624 | -0.4730417911734280 | 1.5764966889281600 |
| **KIM-1** | pT19 | 48 h | 98.7 | 163.9 | 112.4 | 0.7316938646430610 | 0.1875200457645290 |
| **KIM-1** | pT23 | 48 h | 44.66666667 | 55.66666667 | 54.66666667 | 0.3176151019950050 | 0.29146281414061700 |
| **KIM-1** | pT6 | 48 h | 25.15814909 | 4.08288419 | 37.58527373 | -2.6233652340471700 | 0.579141725270205 |
| **MCP-1** | pT10 | 24 h | 86.7386692 | 46.94342759 | 259.2677972 | -0.8857521240572490 | 1.579695811178990 |
| **MCP-1** | pT23 | 24 h | 119.8873472 | 154.0105539 | 157.962316 | 0.36134981196377800 | 0.3979010193806720 |
| **MCP-1** | pT6 | 24 h | 487.2562088 | 29.89085529 | 455.6387822 | -4.026904485297040 | -0.09679002009769670 |
| **MCP-1** | pT10 | 48 h | 109.9416583 | 39.09433574 | 241.2296193 | -1.491706645409590 | 1.1336689134758400 |
| **MCP-1** | pT19 | 48 h | 226 | 452.4 | 223.5 | 1.0012761566872700 | -0.01604794123186990 |
| **MCP-1** | pT23 | 48 h | 178.7149029 | 219.4653836 | 168.9673434 | 0.2963334559549550 | -0.08091550326462300 |
| **MCP-1** | pT6 | 48 h | 336.6751014 | 17.08639445 | 433.6457888 | -4.30043713298143 | 0.3651600678874780 |
| **NGAL** | pT10 | 24 h | 510.426663 | 163.0387644 | 2288.866578 | -1.6464886695880600 | 2.164857770350890 |
| **NGAL** | pT23 | 24 h | 214.6216321 | 805.0237943 | 293.3456528 | 1.9072359307252400 | 0.45080611707615 |
| **NGAL** | pT6 | 24 h | 6436.162477 | 97.09423352 | 1429.52316 | -6.050671320517290 | -2.170666752919590 |
| **NGAL** | pT10 | 48 h | 508.6821322 | 115.3035987 | 1516.18513 | -2.1413268783235700 | 1.5756095963653500 |
| **NGAL** | pT19 | 48 h | 2.2 | 1.5 | 3.1 | -0.5525410230287790 | 0.4947646917495780 |
| **NGAL** | pT23 | 48 h | 272.6109175 | 290.2273767 | 302.8582008 | 0.09034027278301890 | 0.151799137008116 |
| **NGAL** | pT6 | 48 h | 4320.025868 | 82.46213228 | 1141.359024 | -5.711164374477080 | -1.9202872763933800 |
| **OPN** | pT10 | 24 h | 255.0493459 | 41.56957863 | 110.0220605 | -2.6171763715294400 | -1.2129835735699200 |
| **OPN** | pT23 | 24 h | 302.1972719 | 311.5482516 | 331.592669 | 0.04396498399796000 | 0.13392147487021600 |
| **OPN** | pT6 | 24 h | 22.00408127 | 17.35011983 | 368.402785 | -0.3428255096364180 | 4.065440926078760 |
| **OPN** | pT10 | 48 h | 218.2509729 | 26.85129798 | 209.2129259 | -3.0229243507319500 | -0.06101609594756210 |
| **OPN** | pT19 | 48 h | 1269.5 | 8708.9 | 1164.3 | 2.778230112040380 | -0.12479755602826400 |
| **OPN** | pT23 | 48 h | 985.8969079 | 1276.65192 | 905.0356489 | 0.3728565253101330 | -0.12346217611526600 |
| **OPN** | pT6 | 48 h | NA | 23.88187821 | 330.9941265 | NA | NA |

16. Table S3. **Biomarker-level paired MSD summary statistics for Figure 3.** (A) Donor-level |$\log_2$FC| values were summarized within each biomarker, time point, and drug as n, mean, SD, and median. These values correspond to the bars and donor-level points shown in Figure 3. (B) For the primary combined analysis, the 24 h and 48 h means were averaged within each biomarker before paired comparison of CsA and VCS across the eight biomarkers. Endpoint-specific analyses compared CsA and VCS across the eight biomarker-level means at 24 h and 48 h. Directionality indicates the number of biomarkers for which CsA exceeded VCS.

| Analysis | n biomarkers | Mean CsA \|log2FC\| | Mean VCS \|log2FC\| | Mean CsA - VCS difference | 95% CI | paired t-test P | Wilcoxon signed-rank P | Directionality | Sign-test P, two-sided |
|---|---|---|---|---|---|---|---|---|---|
| **24/48 h averaged within biomarker** | 8 | 1.667 | 0.948 | 0.720 | 0.373 to 1.066 | 0.0017 | 0.0078 | CsA > VCS in 8/8 biomarkers | 0.0078 |
| **24 h only** | 8 | 1.747 | 1.137 | 0.610 | -0.009 to 1.229 | 0.0526 | 0.0547 | CsA > VCS in 7/8 biomarkers | 0.0703 |
| **48 h only** | 8 | 1.588 | 0.758 | 0.829 | 0.318 to 1.341 | 0.0064 | 0.0078 | CsA > VCS in 8/8 biomarkers | 0.0078 |

17. Figure S2. Mitochondrial Morphology of 2D-cultured PTECs Under Normoxic (A) and Hypoxic (B) Conditions Following CsA and VCS Treatments. PTECs (pT3) were treated for 48 hours with CNI-containing media replenished every 24 hours under either normoxic (21% $O_2$, top panels) or hypoxic (1% $O_2$, bottom panels) conditions. Cells were treated with CsA or VCS at 1, 5, and 10 µM. Mitochondria (green) were stained with Polybrene, and nuclei (blue) were counterstained with Hoechst. No notable differences in PTEC mitochondrial morphology was observed between hypoxic and normoxic conditions. Scale bars = 20 µm.

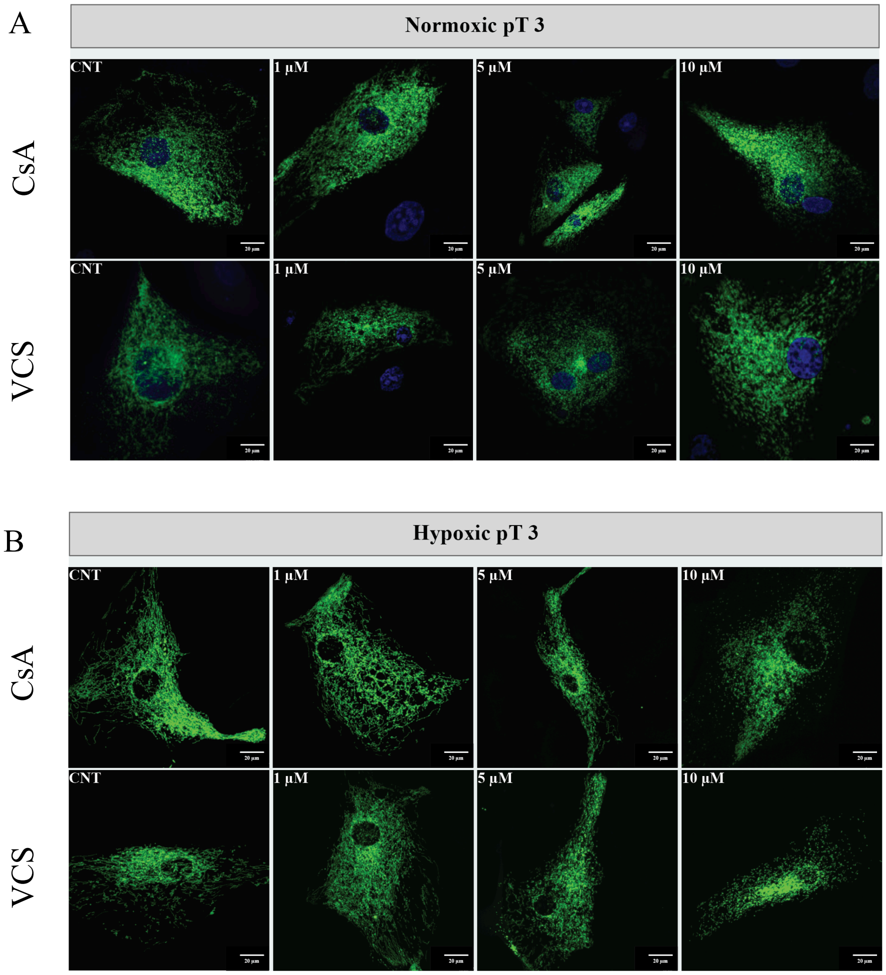

18. Figure S3. Mitochondrial footprint and network branching in 2D-cultured PTECs under normoxic and hypoxic conditions following CsA or VCS treatment. PTECs (pT3) were treated for 48 hours with CsA or VCS at 1, 5, and 10 µM under normoxic (21% $O_2$) or hypoxic (1% $O_2$) conditions, with CNI-containing media replenished every 24 hours. (A) Comparisons of CsA-treated cells vs. controls (CNT) and (B) VCS-treated cells vs. controls show no statistically significant differences in mitochondrial footprint (right panels) or network branching (left panels) under either oxygen condition (One-way ANOVA with Dunnett correction).

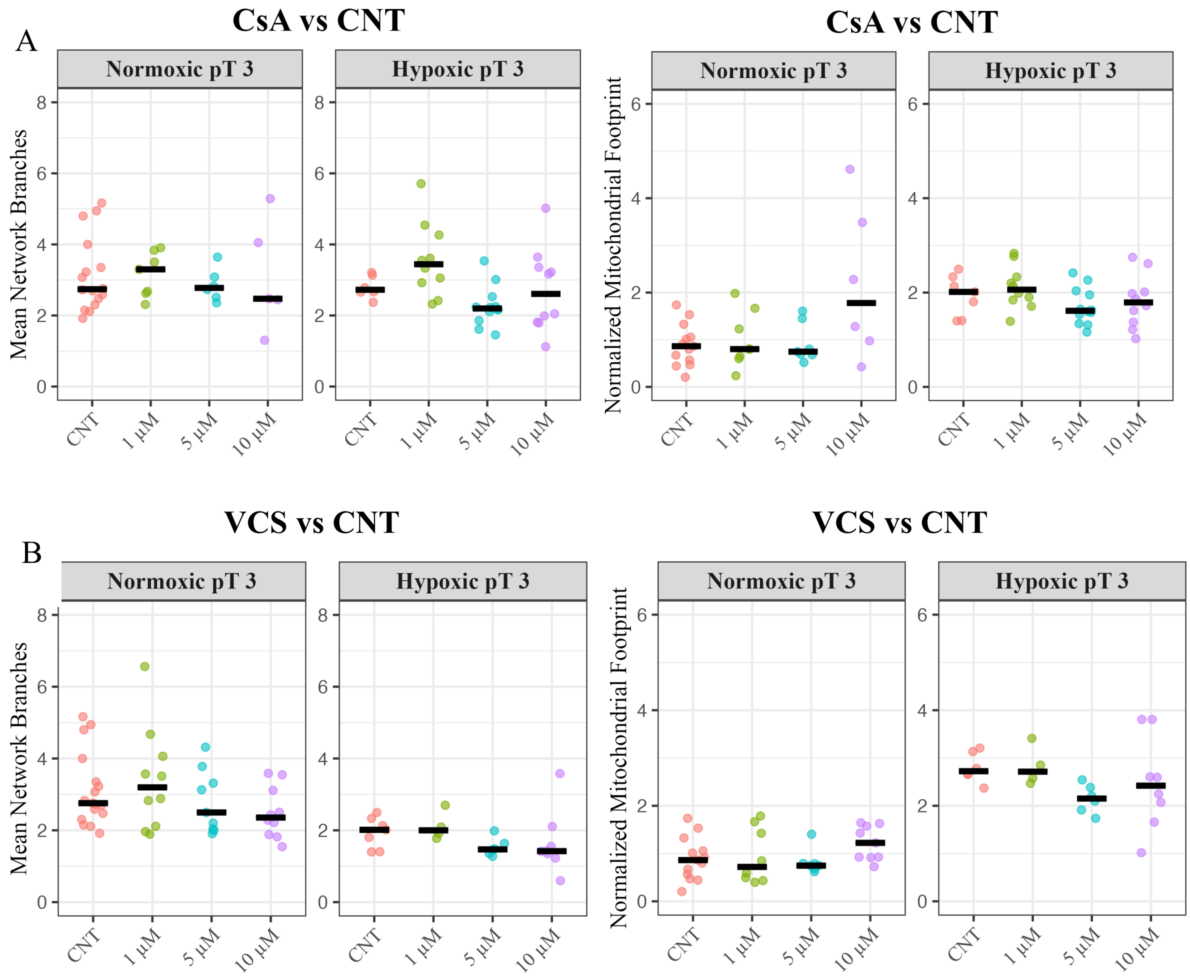

19. Table S4. Summary of RNA-seq Results for CsA and VCS

Summary of RNA-seq Results for CsA and VOC

| Category | CsA | VOC |
|---|---|---|
| **Differentially Expressed Genes** | 9.71% | 3.18% |
| **Significantly Impacted Pathways** | 41 | 39 |
| **Gene Ontology** | 1134 | 1024 |
| **miRNAs** | 144 | 5 |
| **Upstream Regulator Genes** | 268 | 334 |
| **Chemical Upstream Regulators** | 572 | 598 |
| **Significantly Enriched Diseases** | 22 | 35 |

P-value no correction.

20. Table S5. Pathway enrichment analysis for CsA treatments

Pathway Enrichment Calculations for CsA

| Pathway | Observed Genes | Total Pathway Genes | Expected Genes | Fold Enrichment |
|---|---|---|---|---|
| Antigen processing and presentation | 11 | 48 | 4.66 | 2.36 |
| ATP-dependent chromatin remodeling | 11 | 97 | 9.42 | 1.17 |
| Base excision repair | 11 | 41 | 3.98 | 2.76 |
| Cell cycle | 53 | 141 | 13.70 | 3.87 |
| DNA replication | 19 | 32 | 3.11 | 6.11 |
| Ferroptosis | 8 | 35 | 3.40 | 2.35 |
| Gap junction | 13 | 57 | 5.54 | 2.35 |
| Mismatch repair | 8 | 19 | 1.85 | 4.33 |
| Nucleocytoplasmic transport | 22 | 96 | 9.33 | 2.36 |
| Oocyte meiosis | 28 | 112 | 10.88 | 2.57 |
| Progesterone-mediated oocyte maturation | 23 | 93 | 9.03 | 2.55 |
| Protein export | 7 | 29 | 2.82 | 2.48 |
| Protein processing in endoplasmic reticulum | 47 | 157 | 15.25 | 3.08 |
| Small cell lung cancer | 17 | 73 | 7.09 | 2.40 |
| Thyroid hormone synthesis | 10 | 44 | 4.27 | 2.34 |

CsA treatment significantly enriched pathways related to cell cycle regulation, DNA replication, base excision repair, mismatch repair, and ER protein processing. Fold enrichment values reflect the degree to which these pathways were impacted. FDR-adjusted p-value $< 0.05$.

21. Table S6. Pathway enrichment analysis for VCS treatments

Pathway Enrichment Calculations for VOC

| Pathway | Observed Genes | Total Pathway Genes | Expected Genes | Fold Enrichment |
|---|---|---|---|---|
| Antigen processing and presentation | 7 | 48 | 1.15 | 6.10 |
| ATP-dependent chromatin remodeling | 11 | 97 | 2.32 | 4.75 |
| Base excision repair | 8 | 41 | 0.98 | 8.17 |
| Cell cycle | 21 | 141 | 3.37 | 6.23 |
| DNA replication | 11 | 32 | 0.76 | 14.39 |
| Ferroptosis | 4 | 35 | 0.84 | 4.78 |
| Gap junction | 5 | 57 | 1.36 | 3.67 |
| Mismatch repair | 5 | 19 | 0.45 | 11.01 |
| Nucleocytoplasmic transport | 6 | 98 | 2.34 | 2.56 |
| Oocyte meiosis | 10 | 112 | 2.68 | 3.74 |
| Progesterone-mediated oocyte maturation | 12 | 93 | 2.22 | 5.40 |
| Protein export | 6 | 29 | 0.69 | 8.66 |
| Protein processing in endoplasmic reticulum | 34 | 157 | 3.75 | 9.06 |
| Small cell lung cancer | 3 | 73 | 1.74 | 1.72 |
| Thyroid hormone synthesis | 8 | 44 | 1.05 | 7.61 |

VCS treatment demonstrated targeted enrichment in DNA replication, mismatch repair, and protein processing in the ER, with higher fold enrichment values for thyroid hormone synthesis and base excision repair compared to CsA. FDR-adjusted p-value $< 0.05$.

22. Figure S4. Cell cycle pathway analysis in PTECs treated with CsA. KEGG Pathway analysis of the cell cycle in PTECs treated with CsA show genes with significant upregulation (Log2 fold change > 0) highlighted in red, while genes with significant downregulation (Log2 fold change < 0) are shown in blue. This overall pattern suggests a shift toward cell cycle arrest or slowed progression through key phases of the cell cycle in response to CsA treatment.

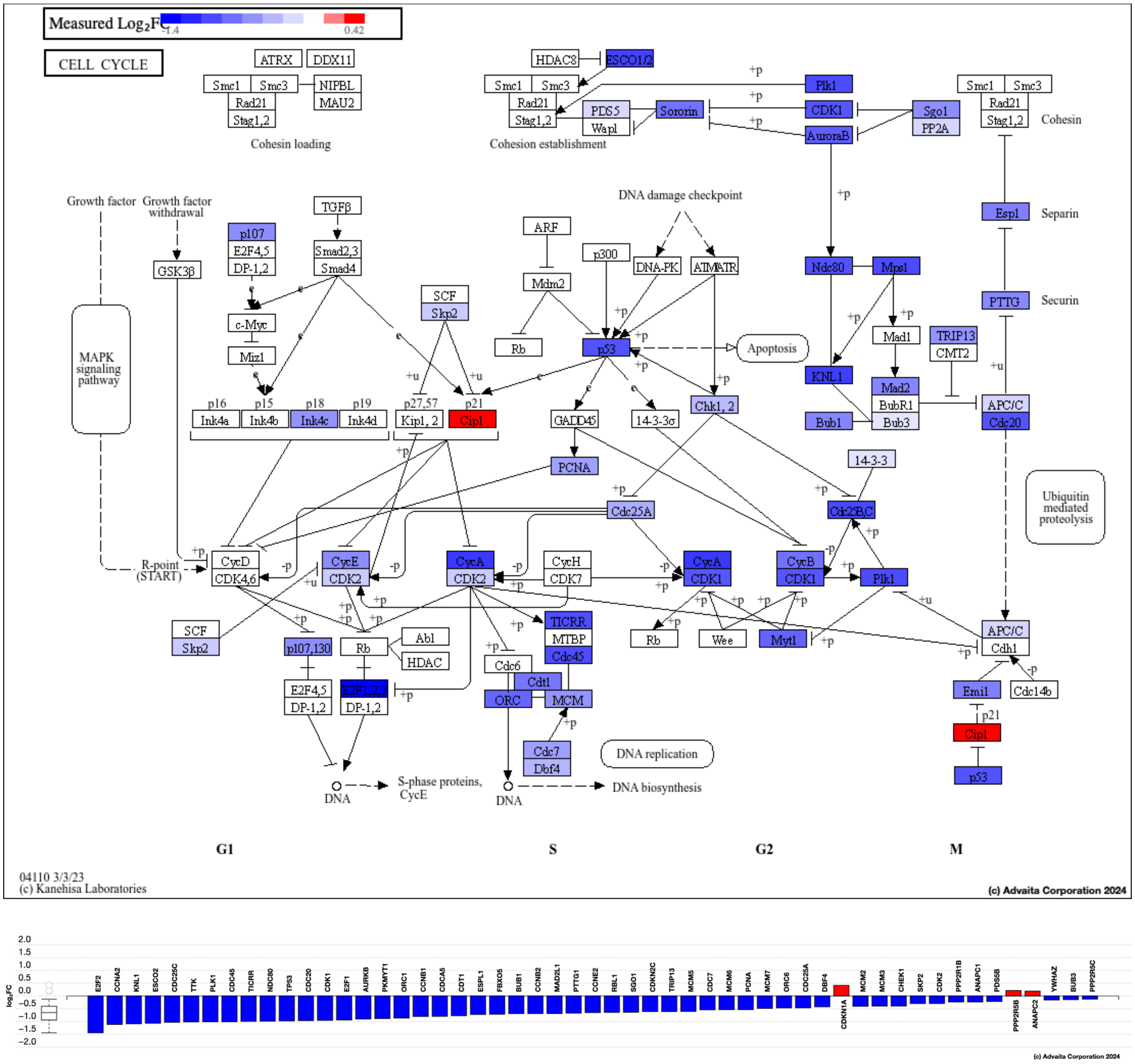

23. Figure S5. Cell cycle pathway analysis in PTECs treated with VCS. KEGG pathway analysis of cell cycle regulation in PTECs treated with VCS show downregulated genes (Log2 fold change < 0) highlighted in blue, while upregulated genes (Log2 fold change > 0) are absent from this analysis. Unlike CsA, VCS did not significantly impact upstream regulators, suggesting milder effects on cell cycle regulation.

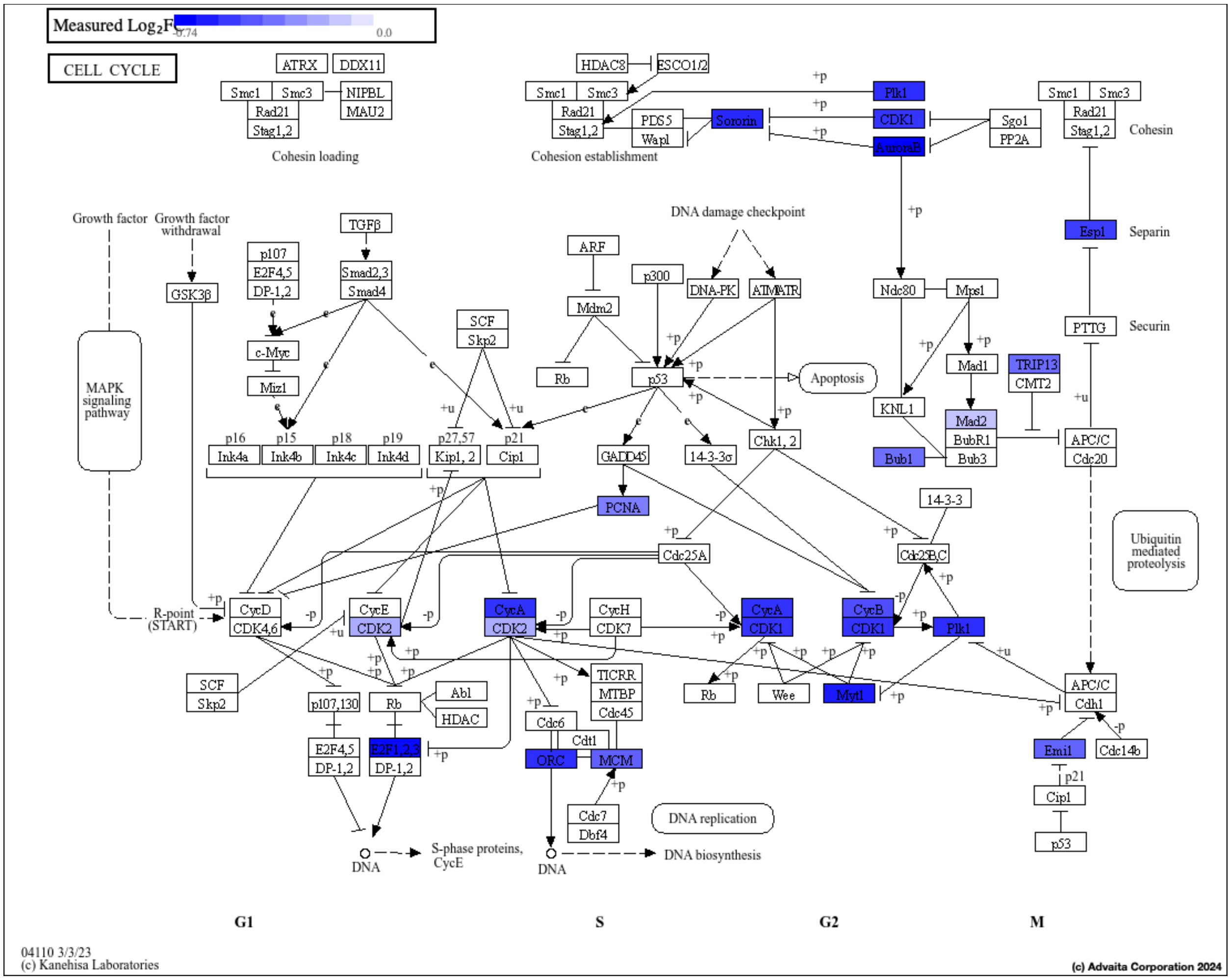


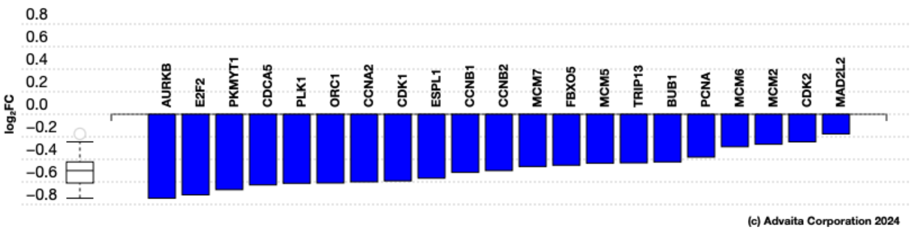

24. Figure S6. ER pathway analysis in PTECs treated with CsA. KEGG pathway analysis of protein processing in the ER following 10 μM CsA treatment of PTECs show increased expression (Log2 fold change > 0) in red, while downregulated genes (Log2 fold change < 0) are in blue. These changes collectively reflect heightened ER stress and disrupted protein homeostasis associated with CsA treatment.

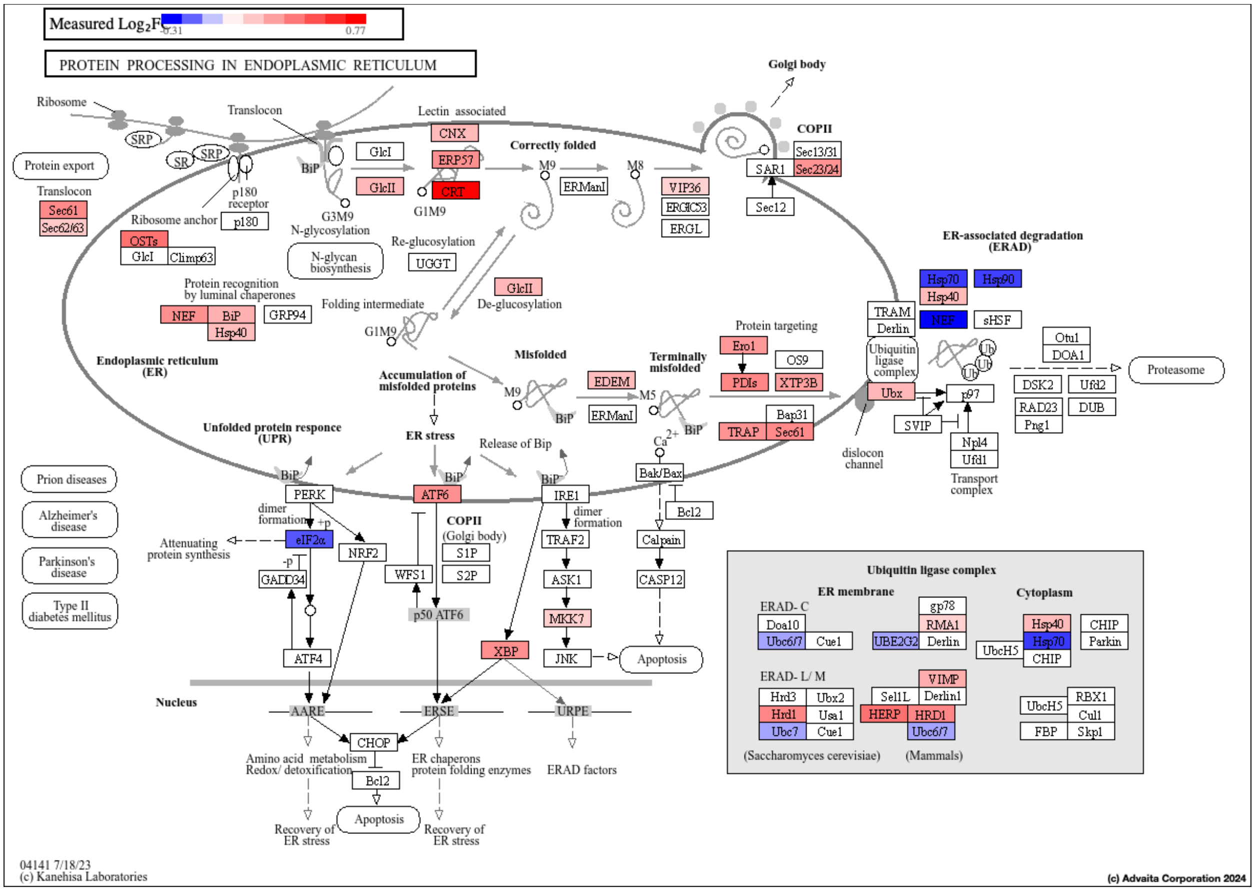


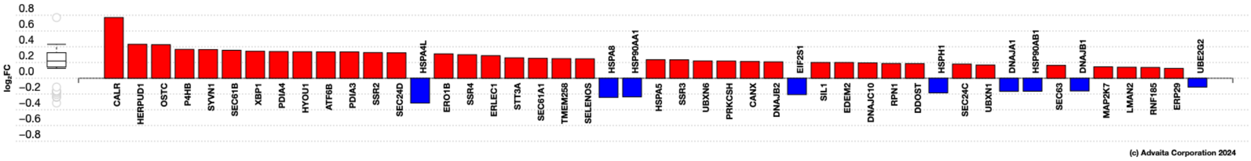

25. Figure S7. ER pathway analysis in PTECs treated with VCS. KEGG pathway analysis of protein processing in the ER following 10 µM VCS treatment of PTECs show increased expression (Log2 fold change > 0) in red, while downregulated genes (Log2 fold change < 0) are in blue. The figure provides insight into the role of VCS in modulating ER proteostasis, with reduced disruption compared to CsA treatment.

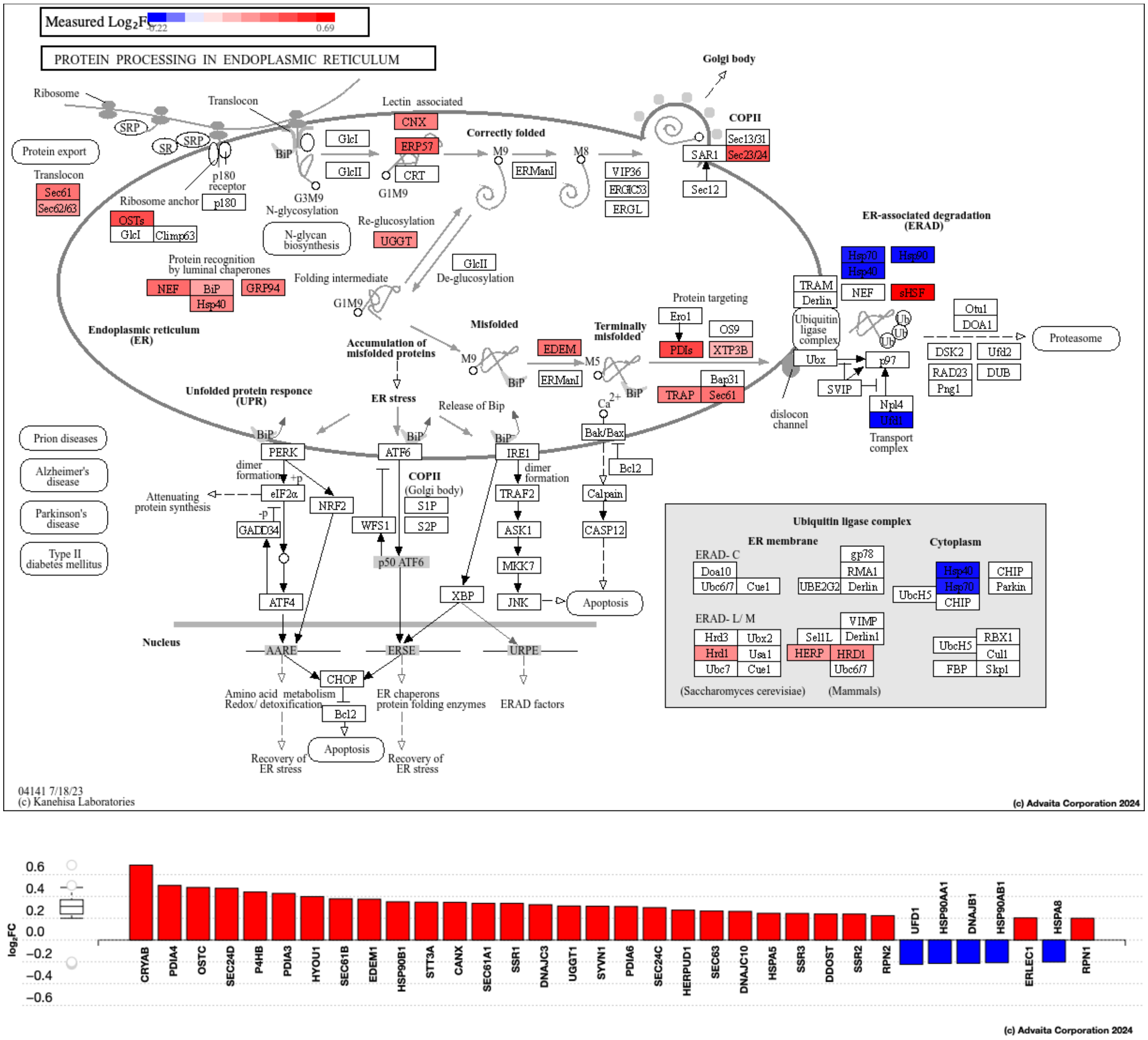

26. Figure S8. Exploratory miRNA-target enrichment analysis based on RNA-seq differential-expression profiles. (A) Venn diagram showing target sets enriched among differentially expressed transcripts after CsA (purple) and VCS (yellow) treatments compared to control. (B) Table showing representative miRNA target sets, the number of differentially expressed target genes, and associated p-values. (FDR-adjusted p-value < 0.05). This analysis infers miRNA-regulatory associations from mRNA target enrichment and does not represent direct miRNA abundance measurements.

A

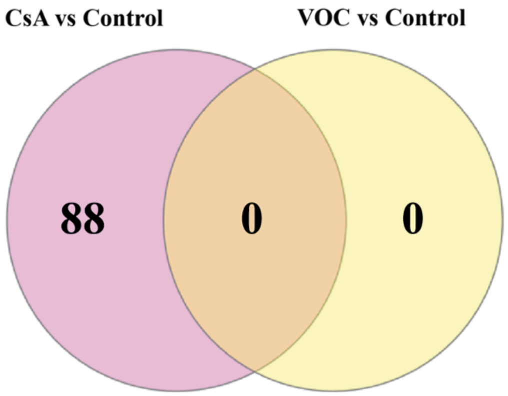


B

| miRNA | VOC vs CNT DE targets | VOC vs CNT p-value | CsA vs CNT DE targets | CsA vs CNT p-value |
|---|---|---|---|---|
| hsa-miR-144-3p | 17/30 | 0.955 | 89/110 | 3.334e-4 |
| hsa-miR-203a-3p | 17/35 | 0.990 | 110/147 | 0.004 |
| hsa-miR-140-3p | 15/28 | 0.990 | 73/97 | 0.004 |
| hsa-miR-6838-5p | 22/48 | 0.990 | 97/132 | 0.004 |
| hsa-miR-195-5p | 22/48 | 0.990 | 97/132 | 0.004 |
| hsa-miR-16-5p | 22/48 | 0.990 | 97/132 | 0.004 |
| hsa-miR-15a-5p | 22/48 | 0.990 | 97/132 | 0.004 |
| hsa-miR-15b-5p | 22/48 | 0.990 | 97/132 | 0.004 |
| hsa-miR-424-5p | 22/48 | 0.990 | 97/132 | 0.004 |
| hsa-miR-497-5p | 22/48 | 0.990 | 97/132 | 0.004 |

27. Figure S9. Mechanism of calcineurin inhibition by CsA and VCS. An antigen-presenting cell (APC) presents peptide antigens in the context of Major Histocompatibility Complex Class II (MHCII) molecules to the T-cell receptor (TCR) on a T-cell (1), while costimulatory signals are provided by B7 (CD80/CD86) on the APC engaging CD28 on the T-cell (2). These two signals are crucial for full T-cell activation. Upon TCR engagement and costimulation, calcium ($Ca^{2+}$) enters the T-cell cytosol through Calcium Release-Activated Calcium (CRAC) channels. The resulting rise in intracellular $Ca^{2+}$ levels activate calcineurin, a $Ca^{2+}$-dependent phosphatase, which dephosphorylates the Nuclear Factor of Activated T-cells (NFAT), allowing NFAT to translocate into the nucleus and IL-2 gene transcription. IL-2 is essential for T-cell proliferation and allograft rejection responses. In addition to their effects on IL-2 transcription, CNIs indirectly influence mitochondrial homeostasis. The elevated $Ca^{2+}$ can be transferred to mitochondria via mitochondria-associated membranes (MAMs). Under normal conditions, the mitochondrial permeability transition pore (mPTP) remains closed, preserving membrane potential and supporting ATP production. Excessive $Ca^{2+}$ and oxidative stress, however, can favor mPTP opening, leading to decreased ATP generation and increased ROS.

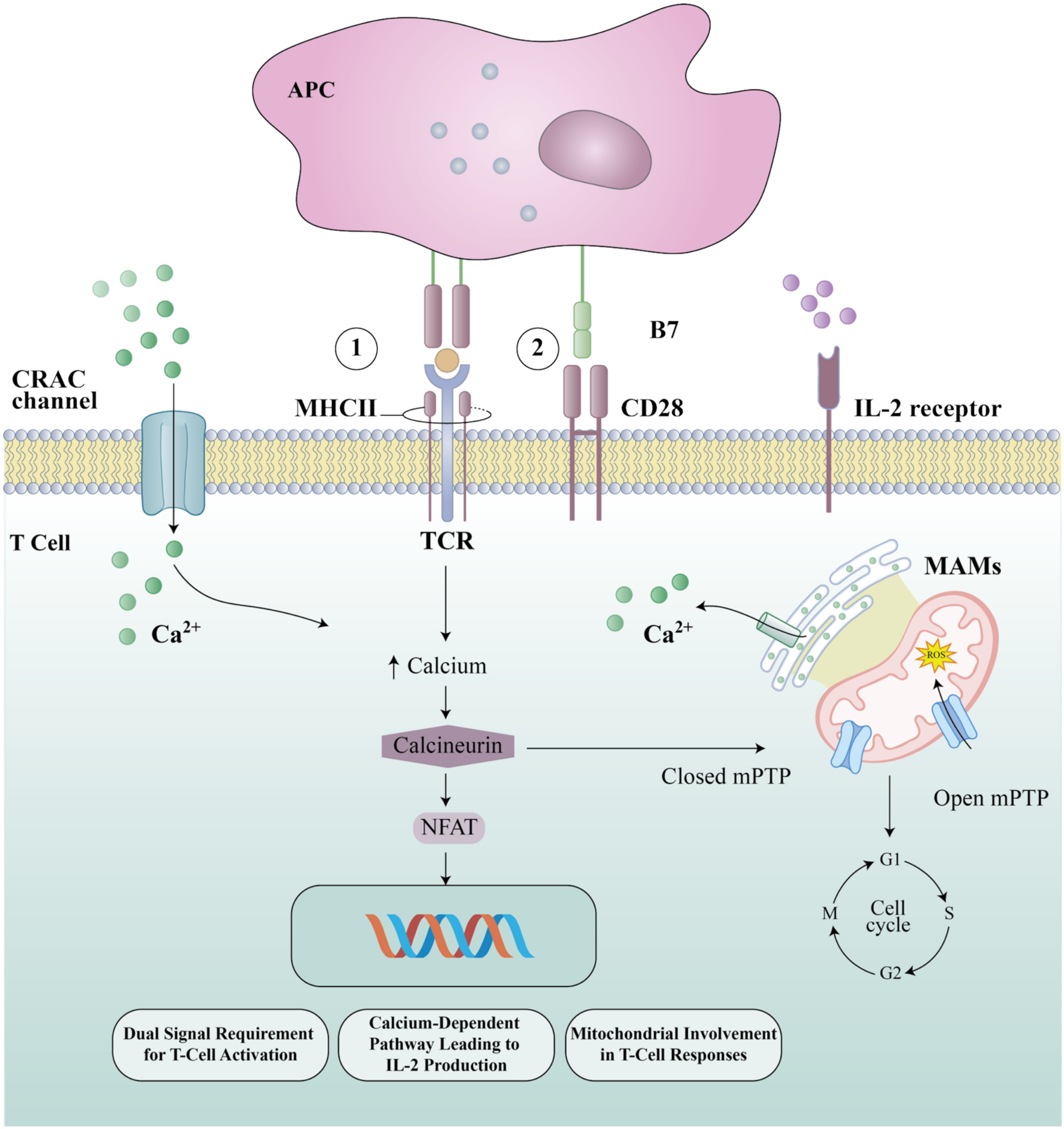
APC
B7
1
2
CRAC channel
MHCII
CD28
IL-2 receptor
T Cell
TCR
MAMs
Ca2+
Ca2+
↑ Calcium
Calcineurin
Closed mPTP
Open mPTP
ROS
NFAT
G1
S
G2
M
Cell cycle
Dual Signal Requirement for T-Cell Activation
Calcium-Dependent Pathway Leading to IL-2 Production
Mitochondrial Involvement in T-Cell Responses
